# A MOUSE ORGANOID PLATFORM FOR MODELING CEREBRAL CORTEX DEVELOPMENT AND CIS-REGULATORY EVOLUTION IN VITRO

**DOI:** 10.1101/2024.09.30.615887

**Authors:** Daniel Medina-Cano, Mohammed T. Islam, Veronika Petrova, Sanjana Dixit, Zerina Balic, Marty G. Yang, Matthias Stadtfeld, Emily S. Wong, Thomas Vierbuchen

**Author notes:** Co-first authors (Daniel Medina-Cano, Mohammed T. Islam, Veronika Petrova). Correspondence (T.V.), (E.S.W.).

## Abstract

Natural selection has shaped the gene regulatory networks that orchestrate the development of the neocortex, leading to diverse neocortical structure and function across mammals, but the molecular and cellular mechanisms driving phenotypic changes have proven difficult to characterize. Here, we develop a reproducible protocol to generate neocortical organoids from mouse epiblast stem cells (EpiSCs) that gives rise to diverse cortical cell types, including distinct classes of excitatory neurons (pre-plate, deep-layer, and upper-layer) and glia (oligodendrocyte precursor cells, myelinating oligodendrocytes, astrocytes, ependymal cells). Cortical organoids develop with similar kinetics to the mouse cortex in vivo and begin to exhibit features of maturation in glia and neuronal cell types relatively rapidly compared to human brain organoids. Using this new protocol, we generated cortical organoids from F1 hybrid EpiSCs derived from crosses between standard laboratory mice (C57BL/6J) and four wild-derived mouse strains from distinct sub-species spanning ∼1M years of evolutionary divergence. This allowed us to comprehensively map cis-acting transcriptional regulatory variation across developing cortical cell types using scRNA-seq. We identify hundreds of genes that exhibit dynamic allelic imbalances during cortical neurogenesis, providing the first insight into the developmental mechanisms underpinning changes in cortical structure and function between mouse strains. These experimental methods and cellular resources represent a powerful new platform for investigating mechanisms of gene regulation in the developing cerebral cortex.

## INTRODUCTION

The mammalian cerebral cortex is comprised of a diverse array of neuronal and glial cell types, which work together to form neural circuits that are critical for higher cognitive processes^1,2^. Generation of these functionally specialized cell types during development requires dynamic and tightly controlled changes in gene expression, which are orchestrated by complex gene regulatory networks^3–6^. Over evolutionary timescales, natural selection has shaped the gene regulatory networks that control neuronal cell fate specification and terminal differentiation, leading to changes in neural circuit formation, neuroanatomy, function, and ultimately behavior across mammals. Determining the mechanisms of gene regulation that govern cerebral cortex development and understanding how genetic variation contributes to variation in cortical structure and function between individuals and between species are central questions in neurodevelopmental biology^7,8^.

Over the last several decades, mice have been the predominant experimental model system used to study the genetic regulation of cerebral cortex development. Genetically engineered mice have played a central role in unraveling how cortical cell types are generated during embryonic development and defining critical transcriptional regulators that establish and refine diverse cortical cell types^6^. More recently, pluripotent stem cell (PSC)-derived cortical organoids have emerged as a powerful, reduced complexity model system that can recapitulate early stages of cortical development *in vitro*^9–11^. Importantly, organoids can be generated from human and primate pluripotent stem cells (PSCs), which has made it experimentally tractable to study mechanisms of human embryonic cortical development^5,12–15^. However, although the first neural organoids were made from mouse PSCs, essentially all neural organoid studies over the last decade have used human or primate PSCs^10,16,17^. This is in large part due to the lack of robust and reproducible protocols for generating mouse neural organoids from pluripotent stem cells.

We previously developed a mouse stem cell-derived forebrain organoid that faithfully recapitulates the development of the hippocampus, cortical hem, and prethalamus^18^. Mouse cortical organoids have the potential to be a powerful complement both to *in vivo* studies of cortical development in mice and to human cortical organoid studies. For example, the scalable nature and ease of experimental access of in vitro organoid models can facilitate experimental techniques and designs that are difficult (or in many cases impossible) to perform *in vivo*, such as live imaging and genetic screens^19–22^. Mouse cortical organoids would also make it possible to utilize the extensive available catalog of mouse genetic resources (including genetically-engineered mouse strains such as gene knockouts and reporter alleles, genetically diverse inbred strains, outbred stocks developed for genetic mapping) to study molecular and cellular mechanisms of cortical development^23,24^. Importantly, new findings from mouse cortical organoid models can be rigorously evaluated in vivo, which is difficult/impossible in most cases for findings from human brain organoids. Finally, the relative ease of genetic engineering in mouse PSCs makes it possible to directly perform experiments and evaluate phenotypes in developing cortical cell types without first having to make genetically engineered mice, which can help to increase experimental throughput and to reduce the number of live animals required for a given project.

Here, we develop a new protocol to reproducibly generate cerebral cortical organoids from mouse epiblast stem cells (EpiSCs) that enables in-depth mechanistic studies of cortical development *in vitro*. Cortical organoids develop with similar kinetics to the mouse cortex *in vivo*, sequentially giving rise to pre-plate, deep-layer, and upper-layer neurons. After cortical neurogenesis, cortical radial glial cells (RGCs) generate astrocytes, oligodendrocytes, or ependymal cells, recapitulating the ensemble of RGC-derived cells present in the developing cortex. Importantly, we can also see evidence for the acquisition of postnatal gene expression programs and functional maturation of multiple cell types within cortical organoids over a period of several weeks. For example, cortical neurons begin to accumulate DNA methylation at CpA dinucleotides, astrocytes activate gene expression programs associated with postnatal development *in vivo*, and OPCs differentiate into myelinating oligodendrocytes.

Using this new protocol, we were able to reproducibly generate cortical organoids from a new panel of F1 hybrid iPSC lines derived from crosses between standard laboratory mice (C57BL/6J) and four wild-derived inbred mouse strains from distinct subspecies (n = 16 total iPSC lines). Distinct mouse sub-strains greatly differ in behavior and gross neuroanatomy^25–28^. However, the underlying genetic mechanisms remain poorly characterized. Using scRNA-seq and novel computational tools, we generated a comprehensive map of allele-specific gene expression across developing cortical cell types. We identify hundreds of genes that exhibit dynamic allelic changes during cortical neurogenesis, providing some initial insight into the gene regulatory changes that contribute to changes in cortical structure and function between distinct mouse strains spanning ∼1M years of evolutionary divergence. Taken together, these new experimental methods, cellular resources, and bioinformatic approaches represent a powerful platform for investigating mechanisms of gene regulation during brain development across evolutionary timescales.

## RESULTS

### Development of a mouse organoid model for recapitulating cerebral cortex development

The cerebral cortex is derived from the anterior, dorsolateral region of the developing neural tube. To generate regionalized cerebral cortical organoids, it is necessary to provide the appropriate combination and intensity of signaling cues that control anterior-posterior and dorsal-ventral patterning of the developing neural tube^10,17,29–32^. In the embryo, acquisition of anterior neuroectodermal fates requires potent inhibition of Wnt signaling immediately prior to and during the early stages of neural induction^33,34^. To generate mouse cortical organoids, we dissociated EpiSCs into single cells and formed embryoid bodies (EBs) of ∼1000 EpiSCs in Aggrewell plates in neural induction media (TGF-*β* and BMP inhibition -- dual SMAD inhibition)^18^. In addition to the neural induction media, we added an inhibitor of Wnt secretion (LGK479) to promote the acquisition of anterior neuroectodermal identity (telencephalon) while also inhibiting Shh signaling (via cyclopamine) to prevent the acquisition of ventral telencephalic identity^35^. After 24h, EpiSC-derived EBs are transferred into Matrigel domes to impart apicobasal polarity within the nascent neuroepithelial tissue^36^. However, we observed that culture of organoids in these initial patterning conditions beyond 48 hours led to limited growth and expansion of cortical neuroepithelia and cell death. To overcome this issue, we found that removing TGF-B, BMP, WNT inhibitors after the first 48 hours and adding 1% knockout serum replacement (KSR) on d3-4 improved survival and growth. In addition, a 24h pulse of Wnt signaling (via addition of the GSK-3*β* inhibitor CHIR99021) further improved growth and survival, as has been observed previously in human cortical organoids^37^. After the first four days, organoids are removed from matrigel domes and transferred to low adherence plates in an orbital shaker in basal media containing 20% KSR for long-term culture (**Fig. 1A**).

**Figure 1:**
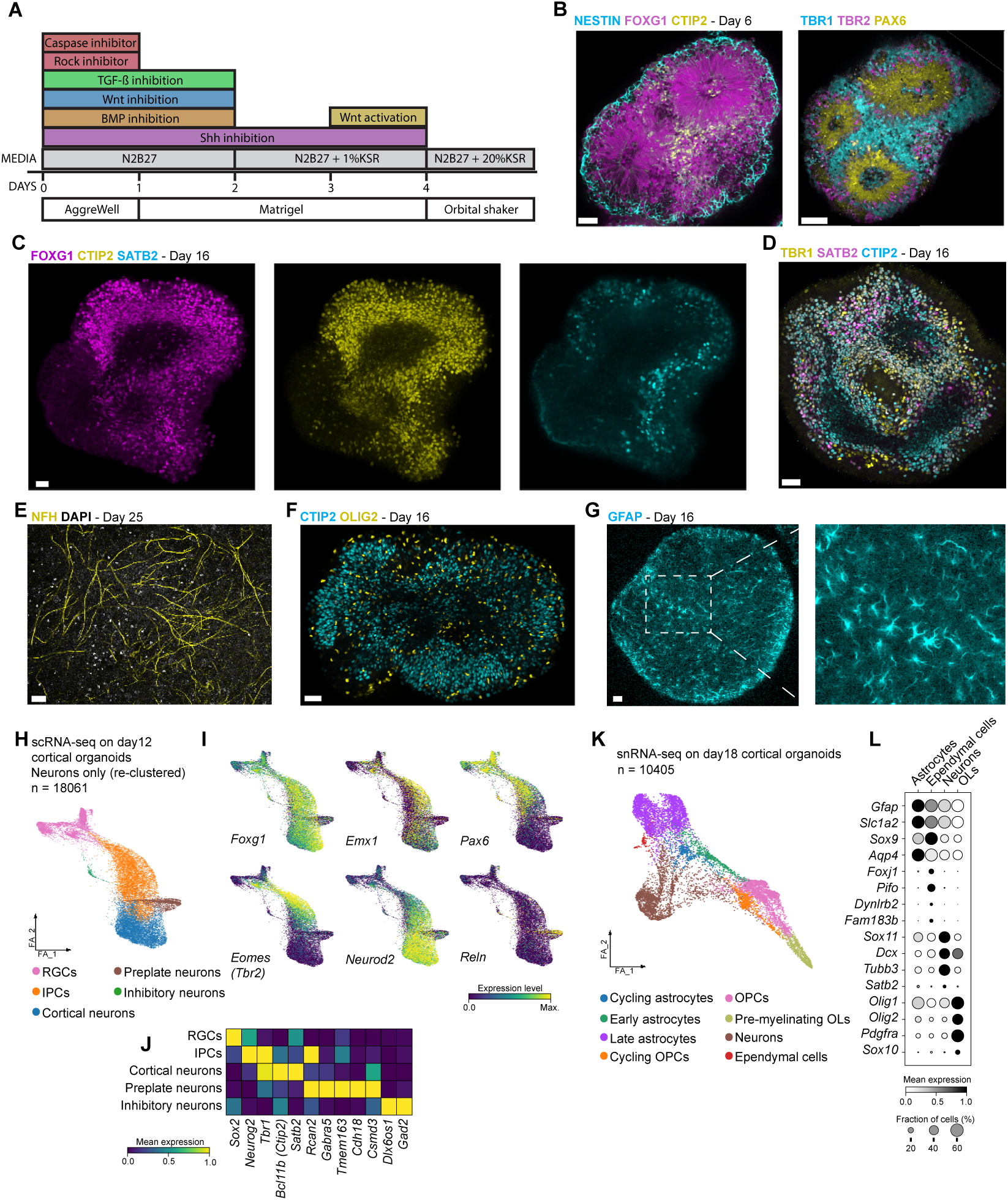
Mouse cortical organoids recapitulate the cellular diversity of the developing cerebral cortex. (**A**) Overview of the protocol for generation of neocortical organoids from mouse epiblast stem cells (EpiSCs). (**B**) Immunostaining of neocortical organoids (d6) for markers of telencephalic progenitors (FOXG1), radial glia (NESTIN), deep-layer cortical neurons (BCL11B/CTIP2, left), and cortical radial glia (PAX6), basal/intermediate progenitors (EOMES/TBR2), and pre-plate/deep-layer cortical neurons (TBR1). (**C**) Immunostaining of neocortical organoids (d16) for FOXG1, BCL11B/ CTIP2, and SATB2 (upper-layer cortical neurons). (**D**) Immunostaining of neocortical organoids (d16) for BCL11B/ CTIP2, TBR1, and SATB2. (**E**) Immunostaining of a d25 organoid for the axonal marker Neurofilament H (NFH). (**F-G**) Immunostaining of neocortical organoids (d16) for markers of cortical neurons (BCL11B/CTIP2), oligodendrocytes (OLIG2), and astrocytes (GFAP). (**H**) Force-directed layout representation of the neuronal cells identified by scRNA-seq in d12 cortical organoids, isolated from the other detected cell types and re-clustered. (**I**) Expression of cell type-specific marker genes (related to A): telencephalon (Foxg1), dorsal telencephalon (Emx1), cortical radial glial cells (Pax6), intermediate progenitors (Eomes/Tbr2), neurons (Neurod2), and preplate neurons (Reln). (**J**) Heatmap of expression of cell type-specific marker genes in selected clusters identified in d12 scRNA-seq data. (**K**) Force-directed layout representation of snRNA-seq data from d18 cortical organoids, and (L) dot plot of selected genes associated with each of the indicated cell types. Scale bars = 40µm. Abbreviations: IPC = intermediate progenitor cell, OPC = Oligodendrocyte progenitor cell, OL = Oligodendrocyte, RGC = radial glial cell.

Using this protocol, we were able to reproducibly obtain cortical organoids exhibiting homogeneous expression of the telencephalic marker FOXG1 and containing PAX6-expressing radial glial progenitor cells (RGCs), consistent with proper patterning into a cerebral cortical identity (**Fig. 1B**). Cortical organoids undergo apicobasal polarization and form rosette structures that mimic the cytoarchitecture of the early developing cortical neuroepithelium. By d6, RGCs within the organoid have started to generate intermediate progenitor cells (IPCs, TBR2+) and early-born excitatory projection neurons expressing markers characteristic of pre-plate neurons (TBR1) and/or deep-layer projection neurons (TBR1, BCL11B/CTIP2) (**Fig. 1B**). Neurons expressing markers of upper-layer/superficial-layer cortical neurons (SATB2), which are produced after the deep-layer neurons in vivo, are first observed at ∼d9 and are found throughout cortical organoids by d16 (**Fig. 1C,D**). These cortical neurons form long axons that traverse the organoids at later timepoints (d25; **Fig. 1E**). In the developing cortex, RGCs gradually lose their neurogenic potential and progress to generate astrocytes or oligodendrocyte precursor cells (OPCs), between ∼E14.5-E17.5^5^. We observe a similar transition in cortical organoids, with astrocytes (GFAP+) and OPCs (OLIG2+) initially emerging at ∼d10 and present throughout the organoids by d16 (**Fig. 1F,G**).

To more systematically define the identity and diversity of neural cell types present in cortical organoids, we performed single cell RNA-seq (scRNA-seq) on d12 cortical organoids (n = 16 independent EpiSC lines, 4 batches, 4 cell lines/batch; additional details and data reported in **Fig. 5, Fig. S2-3, see Methods**) and single nucleus RNA-seq (snRNA-seq) and single nucleus ATAC-seq (snATAC-seq) on d18 cortical organoids (n=4 distinct EpiSC lines, prepared as 2 batches). Consistent with the results from immunofluorescence staining, scRNA-seq data from d12 organoids identified clusters that exhibited expression of markers of cortical RGCs (*Foxg1, Emx1, Pax6*), intermediate progenitors (*Eomes/Tbr2*, *Neurog2*), and distinct types of cortical neurons including pre-plate neurons (*Tbr1*, *Rcan2*, *Gabra5*, *Tmem163*, *Cdh18*, *Csmd3*^38^) and cortical projection neurons (*Neurod2, Tbr1, Reln, Bcl11b/Ctip2, Satb2*) (**Fig. 1H-J**). In addition, we observed a small number (0.9 % of all cells) of inhibitory neurons that express *Bcl11b*, *Dlx6os1*, and *Gad2* which are likely striatal spiny projection neurons which derive from small areas of the organoid which were inappropriately patterned^39^. Clusters corresponding to each of these cell types were also identifiable by scATAC-seq (**Fig. S1A-B;** discussed in additional detail in **Fig. 2-4**). At d18, snRNA-seq data highlighted more extensive glial differentiation, with clusters corresponding to astrocytes, proliferating oligodendrocyte precursor cells, and post-mitotic oligodendrocytes (**Fig. 1K,L**). In addition to these abundant glial cell types, we also identified a small cluster of cells characterized by the expression of ependymal cell markers *Foxj1* and *Pifo*. Ependymal cells arise from late RGCs and populate the ventricular zone, where they act as a barrier from the ventricles^40,41^. Importantly, each of the neuronal and glial cell types were observed across organoids from different batches and cell lines (n =2 batches, n = 4 cell lines) (**Fig. S1C**). Taken together, these data indicate that this protocol can reproducibly generate mouse cerebral cortex organoids from EpiSCs that develop with kinetics that mirror the developing mouse cortex in vivo and generate the major classes of cerebral cortical cell types.

**Figure 2:**
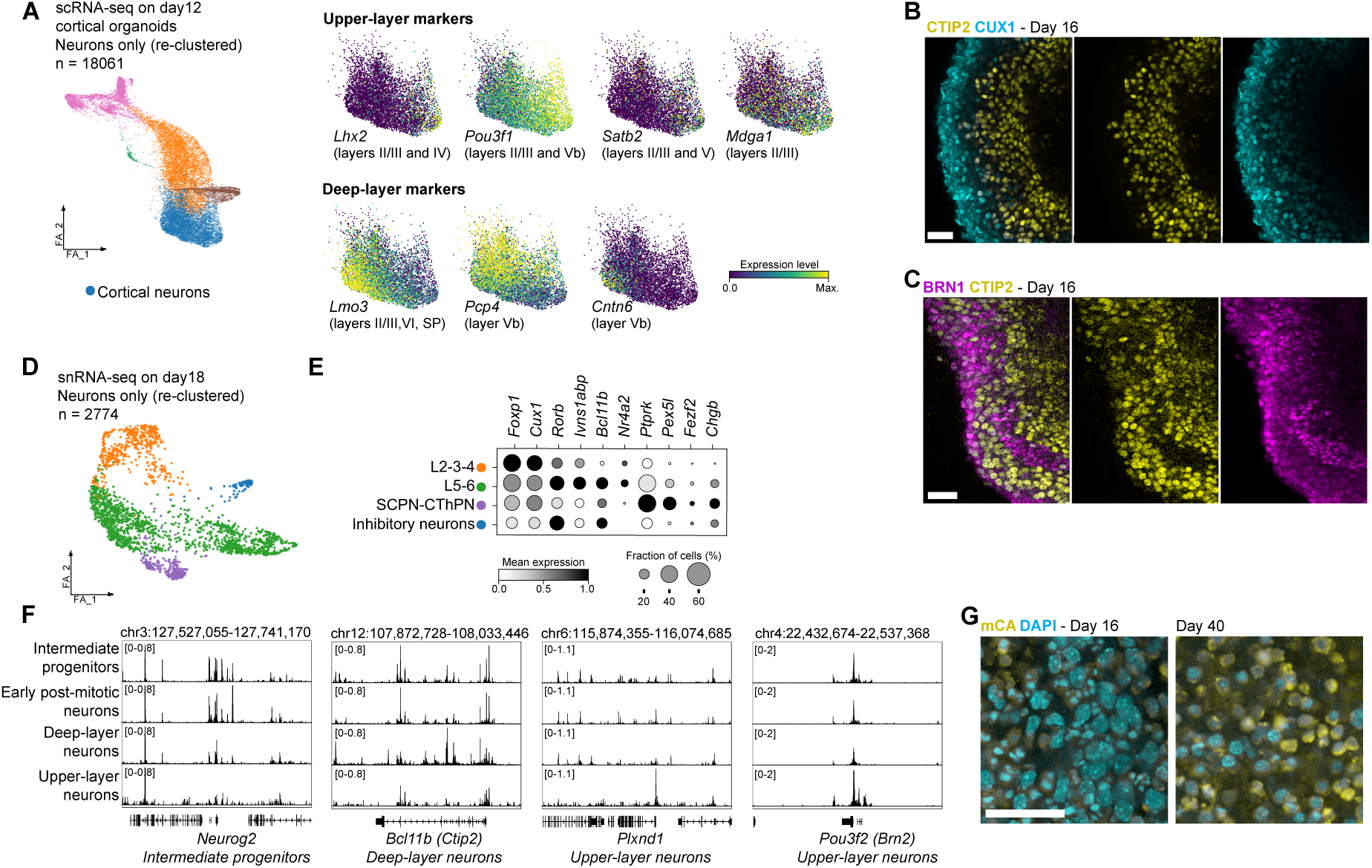
Generation of cortical neuron diversity in mouse cortical organoids. (**A**) Force-directed layout representation of the neuronal cells identified by scRNA-seq in d12 cortical organoids, isolated from the other detected cell types and re-clustered, and expression of selected deep and upper-layer neuron markers in the cortical neuron cluster. (**B-C**) Immunostaining for indicated markers of upper-layer (CUX1, POU3F3/BRN1) and deep-layer (BCL11B/CTIP2). (**D**) Force-directed layout representation of the neuronal cell cluster identified in the d18 snRNA-seq data. The neuronal cluster was isolated from the other detected cell types and re-clustered to identify distinct sub-clusters that might correspond to distinct subtypes of cortical neurons. (**E**) Expression of selected markers of distinct cortical neuron subtypes is plotted for each of the identified cortical neuron sub-clusters. (**F**) Pseudo-bulk chromatin accessibility profiles (snATAC-seq, d18 cortical organoids) at selected cell type-specific gene loci (**G**) Immunostaining for methylated CpA dinucleotides (mCpA) in cortical organoids from d16 and d40. Scale bars = 30µm. Abbreviations: RGC = Radial glial cell, IPC = Intermediate progenitor cell, L2-3-4 = Layers 2-3-4 neurons, SCPN-CThPN = Subcortical projection neuron - Cortico-thalamic projection neuron, L5-6: Layers 5 and 6 neurons.

### Cortical organoids generate major classes of cortical neurons

In the early stages of mouse cortical development (∼E10.5-E11.5), RGCs begin to undergo asymmetric, neurogenic divisions to generate preplate neurons, the earliest-born cortical neurons^42,43^. After preplate neurons, most RGCs proceed to generate deep-layer cortical neurons (layers 5-6) followed by upper-layer cortical neurons (layers 2-4). To further determine the subtype identity of cortical neurons in cortical organoids, we examined expression of a panel of cortical neuron markers characteristic of deep-layer (*Lmo3, Pcp4, Cntn6*) and upper-layer (*Lhx2, Pou3f1, Satb2, Mdga1*) cortical neuron subtypes in d12 scRNA-seq data^4,44^ (**Fig. 2A**). This analysis revealed populations of cortical neurons expressing either deep or upper-layer markers, suggesting that at d12 RGCs in cortical organoids are producing both deep and upper-layer cortical neurons. Consistent with these data, immunofluorescence staining in d16 organoids revealed deep-layer neurons expressing CTIP2+ and upper-layer-like neurons expressing CUX1+ and/or POU3F3/BRN1+, with no/low expression of CTIP2 (**Fig. 2B,C**). Given that more refined transcriptional programs associated with specific subtypes of cortical neurons take days/weeks to be established in vivo, we examined whether further cortical neuron subtype specification in organoids is more apparent at a later timepoint^44^. We revisited the day 18 snRNA-seq data and performed sub-clustering of the “cortical neuron” cluster to see whether subpopulations corresponding to specific subtypes of cortical neurons could be identified (**Fig. 2D,E**). This analysis revealed clusters resembling deep layer neurons (layers 5-6), characterized by expression of *Bcl11b*, as well as upper layer subtypes (layers 2-4), characterized by expression of *Foxp1* and *Cux1* (**Fig. 2E**). These layer-specific identities were also discernable within the scATAC-seq data, with clear differences in chromatin accessibility at cis-regulatory elements in the vicinity of genes critical for deep (*Bcl11b*), and upper-layer (*Pou3f2*/*Brn2*) neuron fate specification (**Fig. 2F**). In addition to the general deep and upper layer clusters, we also observed a smaller cluster of neurons that exhibited expression of some genes characteristic of sub-cerebral and/or cortico-thalamic projection neurons, including *Ptprk*, *Pex5l*, *Chgb*^44^. This suggests the acquisition of some more refined subtype-specific cortical neuron subtypes is occurring in organoids by d18. Collectively, these data indicate cortical organoids recapitulate key features of embryonic cortical neurogenesis, including the generation of pre-plate, deep, and upper-layer cortical neurons.

One advantage of mouse neural organoids compared to human organoids is that mouse neurons undergo post-mitotic neuronal maturation much more rapidly than human neurons (e.g. ∼2-6 weeks in vivo for mice vs. 2-20 years in human cortex^45–47^). To provide initial insight into whether mouse cortical organoids can support the long-term survival and/or maturation of cortical neurons, we cultured mouse cortical organoids for longer periods. In d103 cortical organoids, we observe extensive MAP2+/CTIP2+ neurons (**Fig. S1D,E**), indicating that cortical organoids can survive for extended time periods. To assess whether cortical neurons begin to undergo some cell-intrinsic changes associated with neuronal maturation, we examined the relative abundance of methylated CpA dinucleotides (mCpA), which become abundant in cortical neurons during postnatal development^48,49^. We observe a strong increase in mCpA levels by immunostaining between d16 and d40, consistent with cell-intrinsic maturation occurring in cortical neurons in organoids (**Fig. 2G**). These data suggest that long-term culture of mouse cortical organoids can be a valuable model for studying neuronal maturation in the future.

### Cortical organoids recapitulate development and maturation of oligodendrocytes and astrocytes

During cortical development, RGCs transition from generating neurons to generating glia, including cortical astrocytes and oligodendrocytes^50^. To characterize the differentiation and maturation of glial lineages in cortical organoids, we performed additional analyses to examine oligodendrocyte and astrocyte production in cortical organoids.

Subclustering of oligodendrocyte lineage cells identified in d18 snRNA-seq data identified three clusters. Inspection of the gene expression profiles of cells in each cluster identified a separation between early, *Pdgfra*-expressing, oligodendrocyte progenitor cells (OPCs) and oligodendrocytes that expressed markers associated with differentiation and myelination (*Enpp6*, *Mbp*) (**Fig. 3A**). This continuum of OPC differentiation mirrored transcriptional changes observed in mouse OPCs differentiating into myelinating oligodendrocytes *in vitro* (**Fig. 3B**) (data from^51^). These changes in gene expression paralleled changes in chromatin accessibility at cis-regulatory elements at the loci of genes that play a pivotal role in myelination (e.g. *Myrf, Mag*) (**Fig. 3C**). Immunostaining of d18 cortical organoids confirmed the presence of MBP+ oligodendrocytes, which matured further into MBP+/CNPase+ oligodendrocytes that appeared to be ensheathing axons of cortical neurons (NFH+) (**Fig. 3D,E**). These MBP+ oligodendrocytes persist in older cortical organoids (d30-40) and exhibit characteristic, complex morphologies (**Fig. 3F,G**).

**Figure 3.**
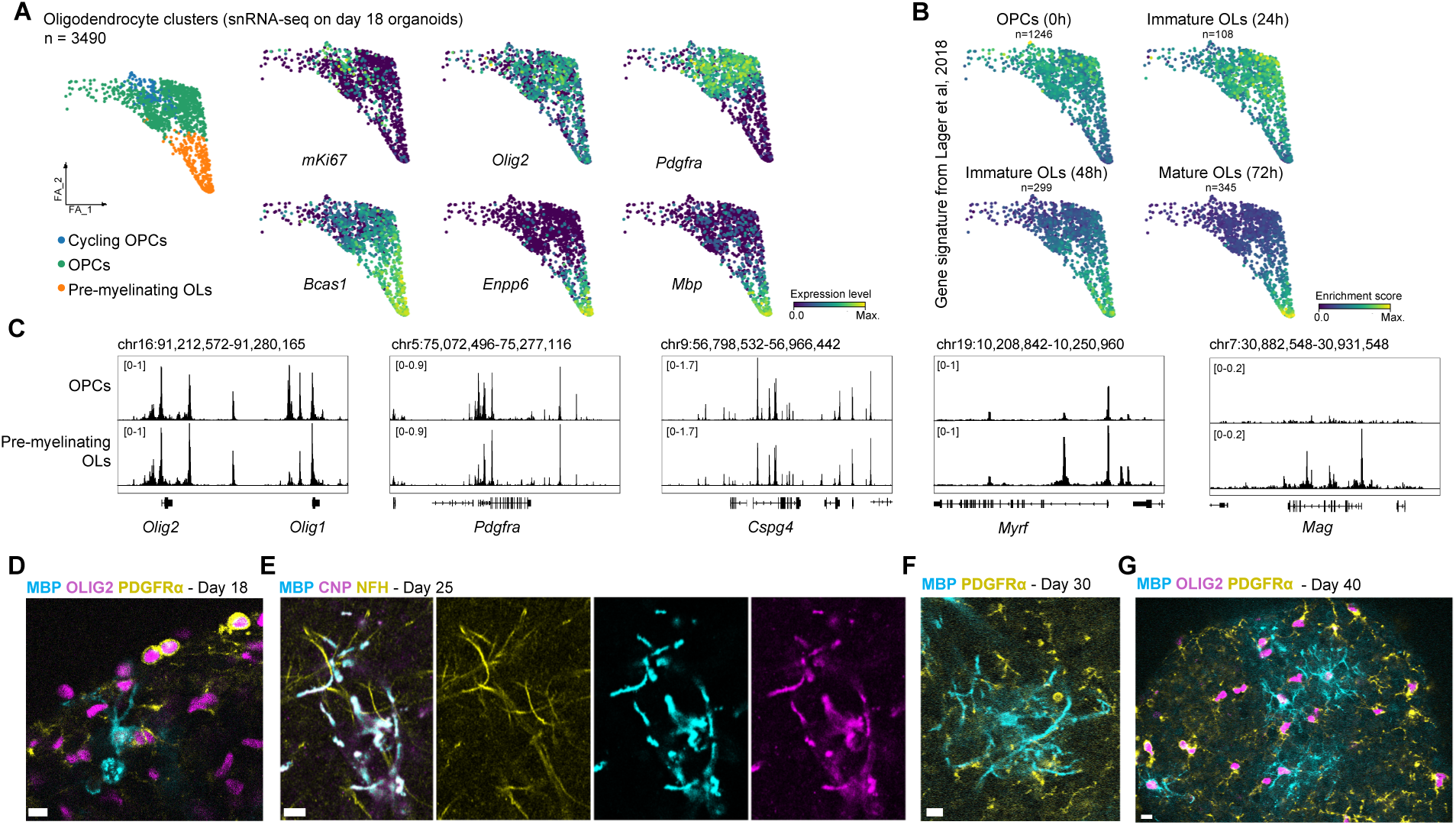
Characterization of oligodendrocyte progenitor cell differentiation and functional maturation. (**A**) Force-directed layout of snRNA-seq data from oligodendrocyte lineage clusters (sub-clustered from d18 organoids). Genes associated with proliferating cells (Mki67), OPCs (Olig2, Pdgfra, Bcas1), and pre-myelinating/myelinating OLs (Enpp6 and Mbp) are highlighted. (**B**) Force-directed layout of snRNA-seq data from oligodendrocyte lineage clusters. Gene enrichment score for genes associated with different stages of OPC/OL differentiation are highlighted. Sample size indicates the number of genes significantly enriched within each cell type, based on gene signatures identified via gene expression profiling of mouse OPC differentiation in vitro (Lager et al, 2018). (**C**) Chromatin accessibility at gene loci associated with oligodendrocyte lineage differentiation: pan-OPC/OL (Olig1/2), OPCs (Pdgfra and Cspg4), and myelinating OLs (Myrf and Mag). Chromatin accessibility profiles were generated from pseudo-bulked scATAC-seq from d18 neocortical organoids. (**D**) Immunostaining of d18 neocortical organoids for markers of OPCs (OLIG2, PDGFRA) and pre-myelinating/ myelinating OLs (MBP). (**E**) Immunostaining of d25 neocortical organoids for markers of myelinating OLs (MBP, CNP) and axons (NFH = neurofilament heavy). (**F**) Immunostaining of d30 neocortical organoids for MBP and PDGFRA. (**G**) Immunostaining of d40 neocortical organoids for OLIG2, MBP, and PDGFRA. Scale bars = 10µm

Next, we performed a similar examination of the development of astrocytes within cortical organoids. Subclustering of cells with astrocyte identity identified from d18 snRNA-seq data revealed sub-populations of astrocytes that appeared to correspond to different degrees of maturation, based on their differential expression of known markers of early/immature (*Egfr*, *Etv4*, and *Grm5*) and more mature (*Clmn*, *Apoe*, *Syne1*, and *Glul*) astrocytes in vivo (**Fig. 4A**)^52^. To evaluate the relative maturation status of astrocytes, we compared snRNA-seq data from astrocytes in d18 cortical organoids to transcriptomic data from purified cortical astrocytes from distinct stages of cortical development (postnatal day 4 [P4] and P60)^52^. This revealed that the “late astrocyte” cluster in cortical organoids has up-regulated a number of genes that are upregulated during cortical astrocyte development in vivo from P4 to P60 (**Fig. 4B**). Similarly, inspection of chromatin accessibility at cis-regulatory elements in the vicinity of mature cortical astrocyte genes, such as *Fam107a* and *Clmn*, indicated increased accessibility at enhancers of these genes that mirror changes that are observed during postnatal development (between P4-P60; **Fig. 4C**). Consistent with this, examination of older organoids (d25) by immunofluorescence staining indicated that many GFAP+ astrocytes also express S100*β*, a marker associated with astrocyte maturation (**Fig. 4D**). At this stage, astrocytes are found throughout the organoids, coating the outer surface and interspersed among the cortical tissue in the interior regions of the organoid. Taken together, these data indicate that RGCs in mouse cortical organoids undergo the expected transition from neurogenesis to gliogenesis, producing oligodendroglia and astrocytes that can undergo further maturation over a period of 1-3 weeks.

**Figure 4.**
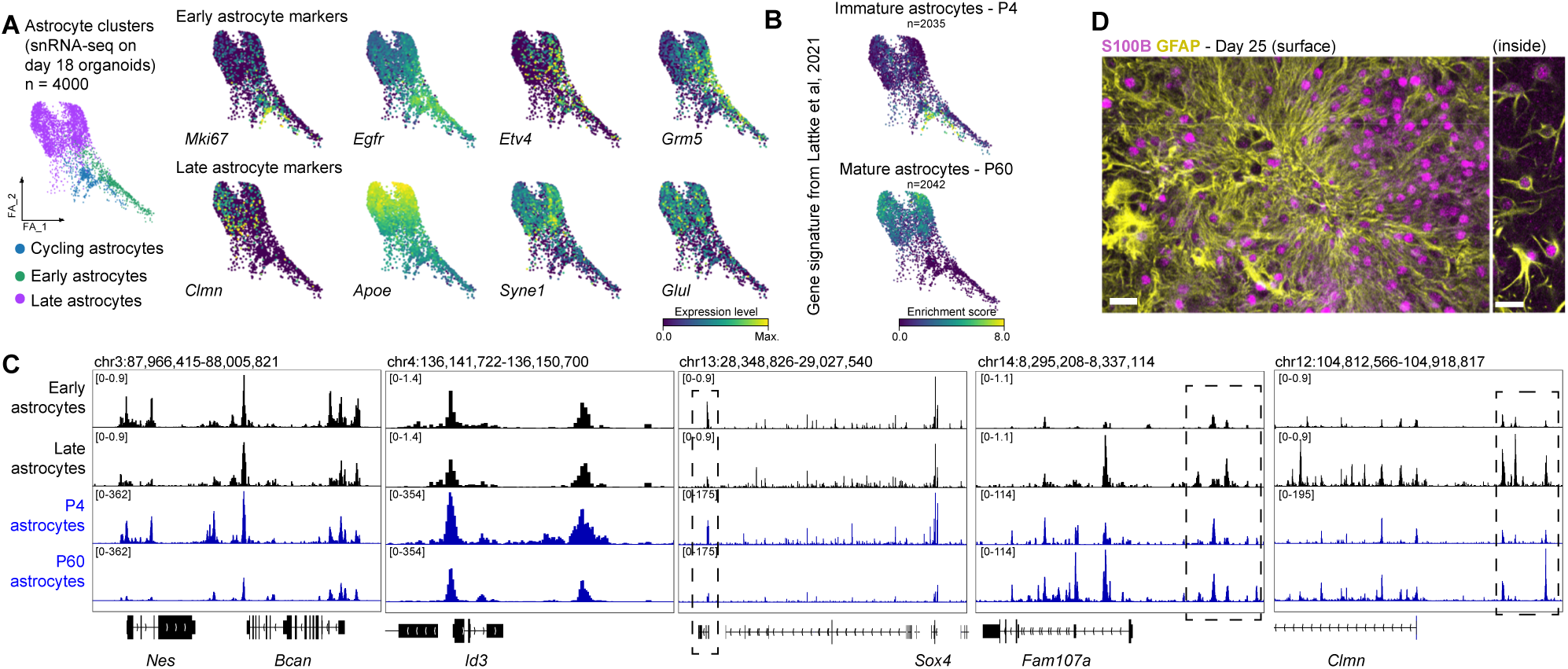
Neocortical astrocytes exhibit transcriptional and epigenomic features of postnatal maturation. (**A**) Force-directed layout representation of the astrocyte clusters identified from snRNA-seq on d18 neocortical organoids. Expression of genes associated with different stages of astrocyte development (identified in15). (**B**) Force-directed layout representation of the astrocyte clusters identified from snRNA-seq on d18 neocortical organoids. Gene enrichment scores for genes associated with immature (P4) or mature (P60) cortical astrocytes are highlighted. Sample size indicates the number of genes significantly enriched within cortical astrocytes from each developmental stage, as identified in Lattke et al, 2021. (C) Chromatin accessibility at genes involved in astrocyte differentiation (Nes, Bcan, Id3, Sox4) and maturation (Fam107a, Clmn). Chromatin accessibility profiles were generated from pseudo-bulked scATAC-seq from d18 neocortical organoids (black) and bulk ATAC-seq profiling of acutely isolated neocortical astrocytes from P4 and P60 mouse brains (Lattke et al, 2021). Dashed rectangles indicate cis-regulatory elements that exhibit changes in accessibility during astrocyte maturation. (D) Immunostaining images of d25 neocortical organoids for the astrocyte markers GFAP and S100. Scale bar = 20µm.

### Generation and characterization of EpiSCs from F1 hybrid crosses between distinct mouse subspecies

Cortical organoids are a powerful model system for mechanistic studies of molecular and cellular features of cortical development. Our new mouse cortical organoid protocol opens up the possibility of performing experiments that take advantage of mouse genetics resources and that would be difficult (or in some cases impossible) to perform in vivo. As a proof-of-concept, we decided to use cortical organoids to characterize how cis-regulatory variation impacts gene expression during cortical development across diverse sub-species of mice.

Commonly used inbred mouse strains, such as C57BL/6, were originally derived from natural populations of *Mus musculus domesticus*^53^. However, there are also a variety of established inbred strains from additional sub-species of mice that exhibit widespread genetic and phenotypic differences compared to C57BL/6J. For example, “wild-derived” inbred strains such as CAST/EiJ (*Mus musculus castaneus),* PWK/PhJ (*Mus musculus musculus),* MOLF/EiJ (*Mus musculus molossinus), and* SPRET/EiJ (*Mus spretus*) span up to ∼1M years of evolutionary divergence from *Mus musculus domesticus* but can still be crossed to standard inbred strains and generate viable (although in several cases infertile) offspring^53–56^. Previous studies have identified extensive neuroanatomical and behavioral differences between these inbred strains. For example, they differ in the total number and relative ratios of neuronal subtypes in several brain regions, they display significant variation in the higher-level organization of axonal connections between brain regions, and they exhibit differences in a variety of behaviors including short-term memory^25–28,57–60^. However, little is known about how the cerebral cortex differs between each of these strains and how genetic changes have contributed to differences in gene regulation during embryonic brain development. This is partially due to the difficulty of working with these wild-derived mouse strains, which can be challenging to consistently obtain, house, and breed^61^.

To specifically evaluate the role of cis-acting regulatory changes in gene expression during cortical development across diverse inbred mouse strains, we chose to generate PSC lines from F1 hybrids between C57Bl/6J females (*Mus musculus domesticus*) and males from four wild-derived inbred strains (CAST/EiJ, MOLF/EiJ, PWK/PhJ, SPRET/EiJ)^54^. This experimental design allows for the comprehensive identification of cis-acting regulatory variation by enabling direct comparison of gene expression levels from the maternal C57BL/6J allele to the paternal allele from each of these wild-derived inbred strains. There is a high frequency of single nucleotide polymorphisms (SNPs) between the C57BL/6J and wild-derived alleles in F1 hybrids (1 SNP every 80-160bp on average), which facilitates allele-specific mapping of many transcripts to either the maternal or paternal alleles using published genome builds from each of the wild-derived inbred strains^55^.

It is hard to derive PSC lines from these crosses using standard techniques, because it is difficult to get wild-derived males from several of these strains to mate consistently with C57BL/6J females^62^. Therefore, we instead chose to generate induced pluripotent stem cell lines (iPSCs) starting from a panel of mouse embryonic fibroblasts (MEFs) from each F1 hybrid cross generated as part of a previous study^54^. For iPSC reprogramming, we used an optimized protocol to generate iPSCs (naïve pluripotent state^63^, see experimental methods). iPSC lines (n = 4-7 clones/F1 genotype) were isolated from individual, nascent iPSC colonies and expanded and banked in the naïve pluripotent state, and then each iPSC line was converted into EpiSCs in vitro, expanded for several passages, and banked in the primed pluripotent state (**Fig. S2A,B;** see methods)^18^. EpiSCs from several of these genetic backgrounds have not been reported previously, so it was not known if standard EpiSC culture conditions (FGF2, Activin A, Wnt pathway inhibitor NVP-TNKS656) would support stable culture of EpiSCs from each of these genetic backgrounds. However, we found that EpiSCs from each of the 4 F1 hybrid backgrounds could be stably maintained in the same culture conditions. Extensive quality control assays, including immunofluorescence staining for pluripotency markers, RNA-seq, and karyotyping were performed to identify high quality EpiSC lines from each genetic background (**Fig. S2C-F;** see **Supplementary table S1** for a summary of quality control assays). In addition, we performed in vitro directed differentiation experiments to evaluate the functional potency of EpiSC lines to differentiate into definitive endoderm and paraxial mesoderm (**Fig. S2G,H**). In summary, we characterized EpiSCs from total of 4 clones from each F1 hybrid genotype (CASTB6 [C57BL/6J X CAST/EiJ], PWKB6 [C57BL/6J X PWK/PhJ], MOLFB6 [C57BL/6J X MOLF/EiJ], SPRETB6 [C57BL/6J X SPRET/EiJ]).

### Generation and characterization of cortical organoids from F1 hybrid EpiSCs

Having established and characterized the F1 hybrid EpiSC panel, we next sought to generate cortical organoids from each EpiSC line to examine cis-regulatory variation during cortical development. We generated cortical organoids from four EpiSC lines from each F1 hybrid background (n = 4 genotypes, n = 4 independent clones/genotype, n = 16 total lines differentiated into organoids in 4 independent batches; **Fig. 5A**, see Methods). Initial characterization by immunostaining confirmed that organoids from each of the F1 EpiSC lines generated CTIP2+ deep-layer cortical neurons and SATB2+ upper-layer cortical neurons (**Fig. S3A**). To examine cell type- and stage-specific gene expression in cortical organoids, we performed scRNA-seq on d12 cortical organoids generated from each F1 EpiSC line (n = 16 total cell lines; initial analysis of neuronal clusters from these data are presented in **Fig. 1H-J**, see Methods). This timepoint was selected because it roughly corresponds to the end of cortical neurogenesis (∼E18.5 in vivo), and thus it should allow for capture of a variety of cortical neuron subtypes at various stages of maturation. UMAP analysis of scRNA-seq data revealed clusters expressing marker genes characteristic of each of the expected classes of cortical cell types, including radial glial progenitors, intermediate progenitors, pre-plate neurons, cortical neurons, immature astrocytes, and OPCs (**Fig. 5B**).

**Figure 5.**
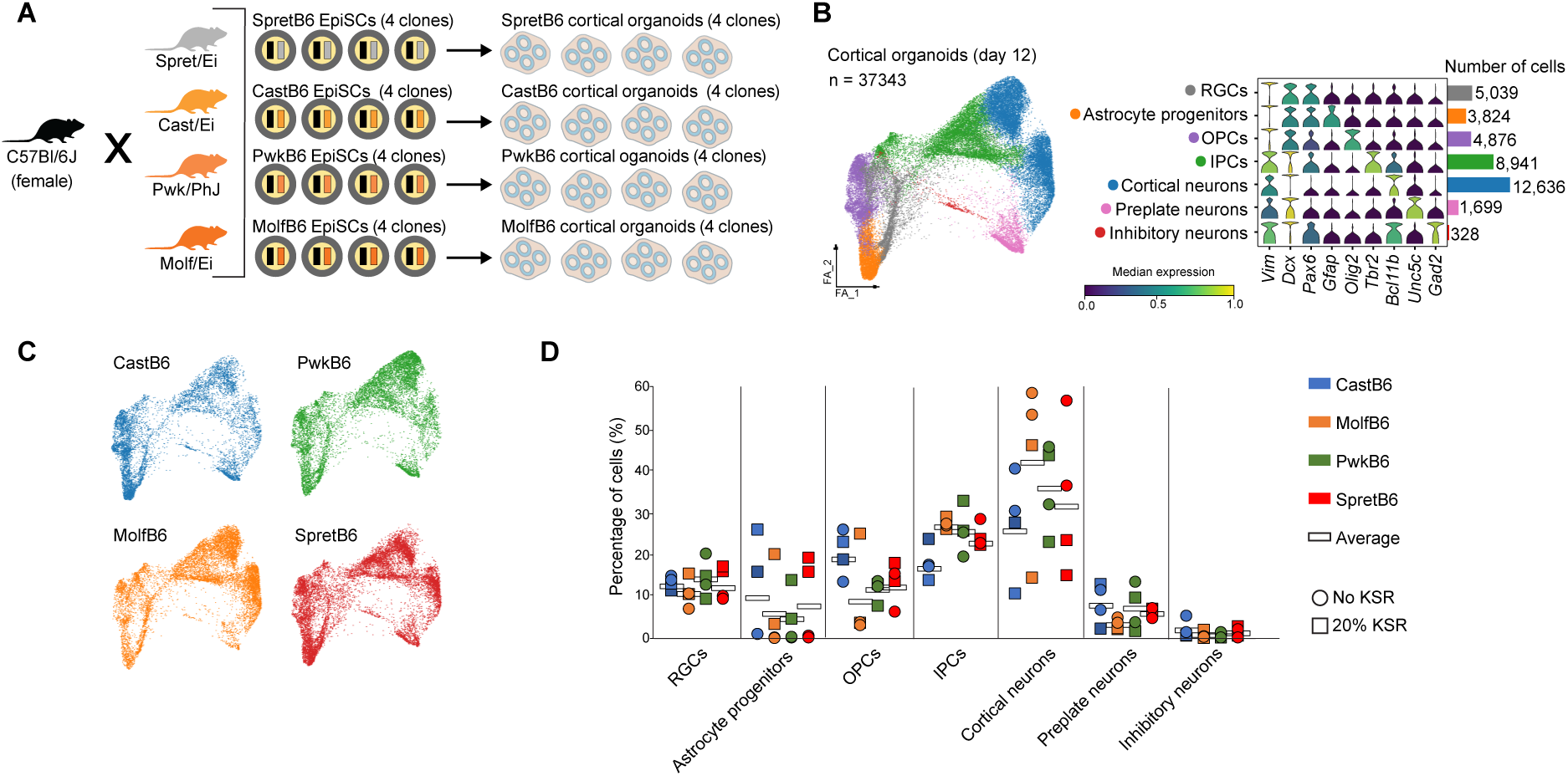
Generation and characterization of neocortical organoids from F1 hybrid EpiSCs. (**A**) Overview of F1 hybrid epiblast stem cell lines (EpiSCs). Four independently derived EpiSC clonal lines from each F1 hybrid genetic background (n =16 EpiSC lines) were used to generate neocortical organoids. (**B**) Force-directed layout representation of scRNA-seq data from d12 neocortical organoids (left panel). Expression of a panel of cell-type specific genes and number of cells of each cell type (right panel). (**C**) Contribution of cells from each F1 hybrid background to the force-directed layout presented in (B). (**D**) Percentage of cells per background and clone on each cell type. Four different backgrounds and four clones per background were used and cultured with or without 20% KSR from day 4. The white rectangle denotes the average between the four clones. Abbreviations: RGC = Radial glial cell, IPC = Intermediate progenitor cell, OPC = Oligodendrocyte progenitor cell

Importantly, each of these cell types are present in cortical organoids from each cell line (n = 16 EpiSC lines), albeit with some quantitative variation in the relative abundance of specific cell types across cell lines (**Fig. 5C,D**). However, given the inherent variability in cell recovery during organoid dissociation and processing for single cell RNA-seq, it is difficult to interpret the significance of these quantitative differences in cell type abundance. One additional source of variability is that two of the four batches of organoids were cultured with 20% knockout serum replacement added to the media after d4, which appears to promote astrocyte differentiation and/or proliferation and causes some differences in gene expression in cortical neurons (**Fig. S3D**). Taken together, these data indicate that each of the F1 hybrid EpiSC lines can be successfully differentiated into cortical organoids, and provide benchmarking data for the performance of the cortical organoid protocol across a genetically diverse panel of EpiSC lines.

### Cell type-specific mapping of cis-regulatory divergence in developing neocortical cell types

Having performed basic characterization of the F1 hybrid cortical organoids from each genetic background, we next sought to further analyze the scRNA-seq data to quantify expression from the C57Bl/6J (B6) and wild-derived inbred strain alleles (Cast/EiJ, Molf/EiJ, Pwk/PhJ and Spret/EiJ). This would allow us to comprehensively assess changes in cis-regulatory control of gene expression in developing cortical cell types across ∼1M years of mouse sub-species evolution. To accomplish this, we mapped exonic reads from scRNA-seq data from d12 cortical organoids from each of the F1 hybrid lines to their respective genomes (C57BL/6J and the appropriate wild-derived inbred strain genome). We generated genomes for the wild-derived inbred strains by incorporating their respective genetic variants into the C57Bl/6J mm10 genome (**Fig. S4A**, see Methods). To increase the sensitivity of read mapping, we incorporated indels as well as SNVs into a hybrid genome for each F1 hybrid line. This increased the percentage of reads that were mappable to each of the two possible genomes by ∼10% compared to SNVs alone, thereby increasing the number of unambiguously aligned reads we could use for calculating allelic ratios (**Fig. S4B**)^64^. To test for differences in expression from each allele, we utilized ASPEN, a newly developed empirical Bayes approach designed for detecting allelic differences using the beta-binomial distribution. The method shrinks gene-wise variance estimates to improve statistical power and to control false discovery rates. To improve accuracy, we also adjusted our statistical test for allelic imbalance to account for a minor empirically derived reference-allele mapping bias (**Fig. S4A,B,** see Methods).

The sparse transcript sampling and 3’ end coverage bias of 10X Genomics scRNA-seq data can lead to less reliable allele-specific measurements for lowly expressed genes^65^. Therefore, to investigate the differences between scRNA-seq results with those obtained from bulk RNA-seq data, we compared scRNA-seq results with matched bulk RNA-seq results from the same cell type. We generated scRNA-seq data from EpiSCs from each of the 4 F1 hybrid strains, and compared allelic quantifications to those generated from bulk RNA-seq (total RNA-seq)(**Fig. 6A**, **Sup. Fig. S4C, see methods**). This revealed a concordant correlation (Pearson ρ ∼ 0.61) between allelic ratios. Roughly 80% of genes detected by aggregated (pseudobulking) scRNA-seq as exhibiting allelic imbalance were also detected in bulk RNA-seq data (FDR < 0.1, ∼5000 genes tested per F1 strain; **Supplementary Table S2**).

**Figure 6.**
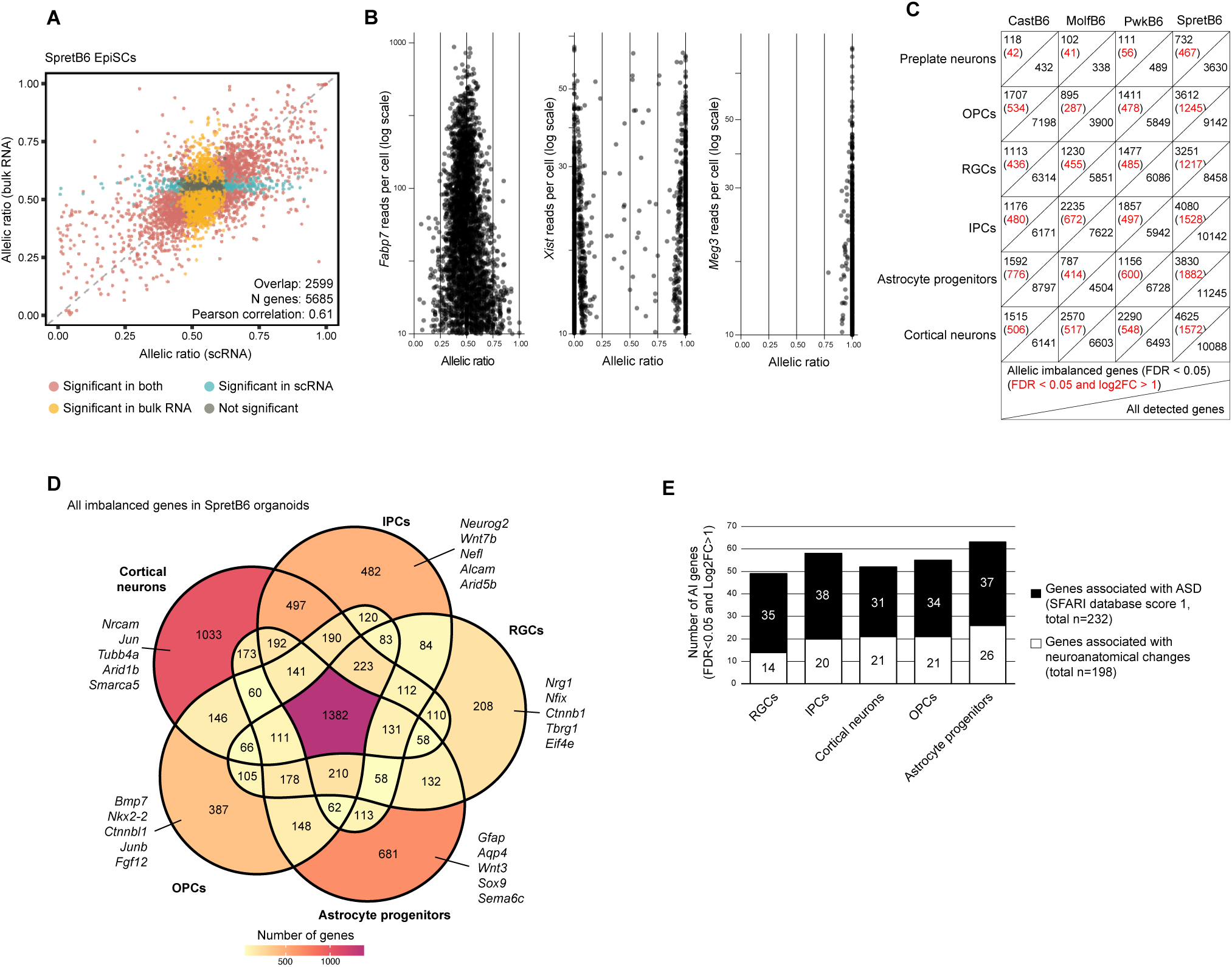
Single-cell allelic mapping in F1-derived cortical organoids identifies gene imbalances associated with neuroanatomical changes. (**A**) Correlation of allelic ratios in SpretB6 EpiSCs as detected by bulk RNA-seq or scRNA-seq. The Pearson coefficient and the number of genes showing overlapping behavior in both datasets are displayed. (**B**) Allelic ratios in single cells, as detected through scRNA-seq on F1-derived cortical organoids, of Fabp7 (MolfB6 organoids), Xist (CastB6 organoids, female cell line), and the imprinted gene Meg3 (MolfB6 organoids). (**C**) Number of genes allelically imbalanced (AI) across cell types and genetic backgrounds as detected by scRNA-seq on day12 cortical organoids. (**D**) Venn diagram of AI genes in SpretB6 organoids and classified by the cell type in which they are detected as imbalanced. (**E**) Histogram representing, out of all the AI genes per cell type – merging all backgrounds – how many have been identified as tightly associated with ASD (SFARI database, score 1 only) or changes within brain neuroanatomy (as identified by Collins et al, 2019).

We next used ASPEN to quantify allelic expression differences in scRNA-seq data from F1 hybrid cortical organoids. The high frequency of SNPs between each allele in these F1 hybrids (∼6-12 times higher frequency than observed between maternal/paternal alleles in an average human) makes it possible to unambiguously assign a higher proportion of reads to each allele than what has been possible in human cells^56,66,67^. For example, we resolved ∼20% of total sequencing reads to alleles in CastB6, MolfB6, and PwkB6 hybrids, and ∼36% in SpretB6. This enabled accurate quantification of allelic ratios for highly expressed genes with single-cell resolution (**Fig. 6B**). We observed clear separation of allelic expression in imprinted and X-linked genes, demonstrating the sensitivity of ASPEN at the single cell level (**Fig. 6B**). For example, we observed strong mono-allelic expression from the maternal (C57BL/6J) allele for the imprinted gene *Meg3* across all cell types examined. Similarly, in female CastB6 organoids, we detected robust X-chromosome inactivation at the single cell level, exemplified by mono-allelic expression of *Xist*.

To quantify allelic imbalances in gene expression in specific cell types (preplate neurons, OPCs, RGCs, IPCs, astrocyte progenitors, cortical neurons), we pseudobulked gene expression data from clusters representing distinct cell types and used these data from each cell type to identify allelic imbalances using ASPEN. Across all cell types, we found that 17-45% of tested genes exhibit significant allelic imbalance (**Fig. 6C, Sup. Fig. S4D**, **Sup. Fig. S5A, and Supplementary tables S3-6**). As expected, the highest frequency of allelic imbalances was observed in comparisons between C57BL/6J and Spret/Ei alleles, consistent with the ∼2-fold higher SNP frequency between these two strains^54,68^. Among all genes that exhibited a significant allelic imbalance in SpretB6 F1 cortical organoids, only 26% (1382/5373) were significantly imbalanced across all cell types examined, meaning that most of the observed genes with significant allelic imbalances were cell type- and/or stage-specific (**Fig. 6D**). Genes that exhibit cell type-specific allelic imbalances included a variety of genes with diverse functions, including transcriptional regulators (Jun, Junb, Neurog2, Nfix, Sox9), chromatin remodelers (*Arid1b, Arid5b, Smarca5*), regulators of cell adhesion (*Nrcam, Alcam*), cytoskeletal components (*Tubb4a, Nefl*), and components of intercellular signaling pathways (*Wnt3*, *Wnt7b, Bmp7, Nrg1*). Additional enrichment analyses (GSEA) on these sets of genes further proved their importance for key aspects of cell type-specific biology (**Sup. Fig. S5B**). For example, GSEA analysis of genes with an allelic imbalance in SpretB6 astrocytes revealed a clear enrichment for terms related to astrocyte morphology, indicating that evolutionary changes to cis-regulation result in strain-specific differences in cell structure and function.

Distinct mouse sub-strains exhibit both tissue-scale neuroanatomical differences and quantitative changes in cell type composition within tissues, including differences in the total and relative abundance of different neuronal cell types^25–28^. To identify changes in cis-regulatory activity that might contribute to these neuroanatomical differences between C57BL/6J mice and the wild-derived inbred strains, we cross-referenced our list of allelic imbalanced genes with a set of genes that cause neuroanatomical phenotypes when knocked-out in mice (n=198)^69^. These analyses identified 30 genes that exhibit allelic imbalance between different inbred strains that exhibit neuroanatomical phenotypes (e.g. *Akt2*, *Id2, Mbd1*) (**Sup. Fig. S5C**). This suggests that differential regulation of these genes might contribute to the observed differences in the adult mouse brain across these inbred strains. Interestingly, we also identified allelic imbalances in many genes (n=56) that have been linked to Intellectual Disability and Autism Spectrum Disorder (ID/ASD) in human populations (SFARI database score = 1, n = 232 genes, **Fig. 6F, Sup. Fig. S5D**). Differential expression of these genes across distinct mouse inbred strains might contribute to the highly variable phenotypic penetrance of ID/ASD mutations when examined in different inbred mouse strains^70,71^.

To understand the extent of the interplay of cis-regulatory variation and cellular context during mammalian brain development, we used generalized linear models to examine how genetic background and cell type may influence observed allelic imbalances. For each gene, we fit models to test whether differences in cell type, genetic background, or a combination of both could explain the observed expression patterns. We imposed constraints on the range of values that the fixed effect parameters could take and evaluated the goodness of fit of the models under different scenarios. For example, to determine if a gene exhibited differential allelic expression between strains and cell types, we compared the fit of the data to two competing models. The null model constrained the strain or cell type terms to zero, while the alternative model allowed the genetic and cell type effect to be fully modeled. Because different cell types naturally have distinct gene expression profiles, a cell type effect may occur simply due to changes in overall gene expression levels. That is, when a gene’s total expression level is high, this can lead to a decrease in allelic imbalance as stochastic effects average out. Hence, we also included total gene expression in our model to account for this.

This analysis indicated that nearly half of the genes (47%) showed differences in allelic expression due to genetic background, regardless of the cell type (340 genes; Bayesian Information Criterion difference >10) (**Supplementary Table 7, Fig. S5E**). This means that genetic differences between strains significantly influence which allele is expressed. We also observed that within the same genetic background, the pattern of allelic expression tends to be more similar across cell types (Spearman ρ ∼ 0.52 – 0.61; **Fig. S5F**). Nevertheless, we also found that cell type can play a substantial role. For 31% of the genes, allelic expression was influenced by both genetic background and cell type, but these effects were independent of each other (219 genes). For 15% of genes, allelic differences were driven solely by the cell type, regardless of the genetic background (109 genes). A smaller group of genes (n= 50; 7%) showed a more complex pattern, where both strain and cell type interacted to modulate allelic imbalance, suggesting that a combination of genetic and cellular factors in these cases influences gene regulation (**Fig. S5G**).

Overall, these results highlight the importance of both genetic differences between the strains and the cell type context in shaping gene expression patterns.

### Dynamics of allelic expression along cortical neurogenesis

We next sought to investigate allelic expression dynamics in developing cortical neurons. We were interested in identifying genes that exhibit allelic imbalance in gene expression in specific stages of cortical neuron differentiation from radial glial cells. Stage-specific allelic imbalances potentially reflect differences in the activity of cell type-specific cis-regulatory element(s) between the C57BL/6 and wild-derived alleles, and thus are thought to be particularly relevant for inter-individual and inter-species variation^72,73^.

To arrange the cells from different samples and genetic backgrounds across the developmental trajectory from radial glial cells to post-mitotic cortical neurons, we performed pseudotemporal analyses using Palantir on d12 scRNA-seq data (**Fig. 7A**, see methods)^74^. Pseudotemporal ordering faithfully captured cortical developmental transitions, based on the overall expression dynamics of known marker genes for each stage of differentiation (**Fig. 7B**). For subsequent analysis of allelic imbalances at the single cell level, we only considered genes that are expressed at a sufficiently high level to be detected across each stage of differentiation so that it is possible to confidently ascertain their allelic imbalance at each stage (3,336 genes across all strains, 1,409 total unique genes, see methods).

**Figure 7.**
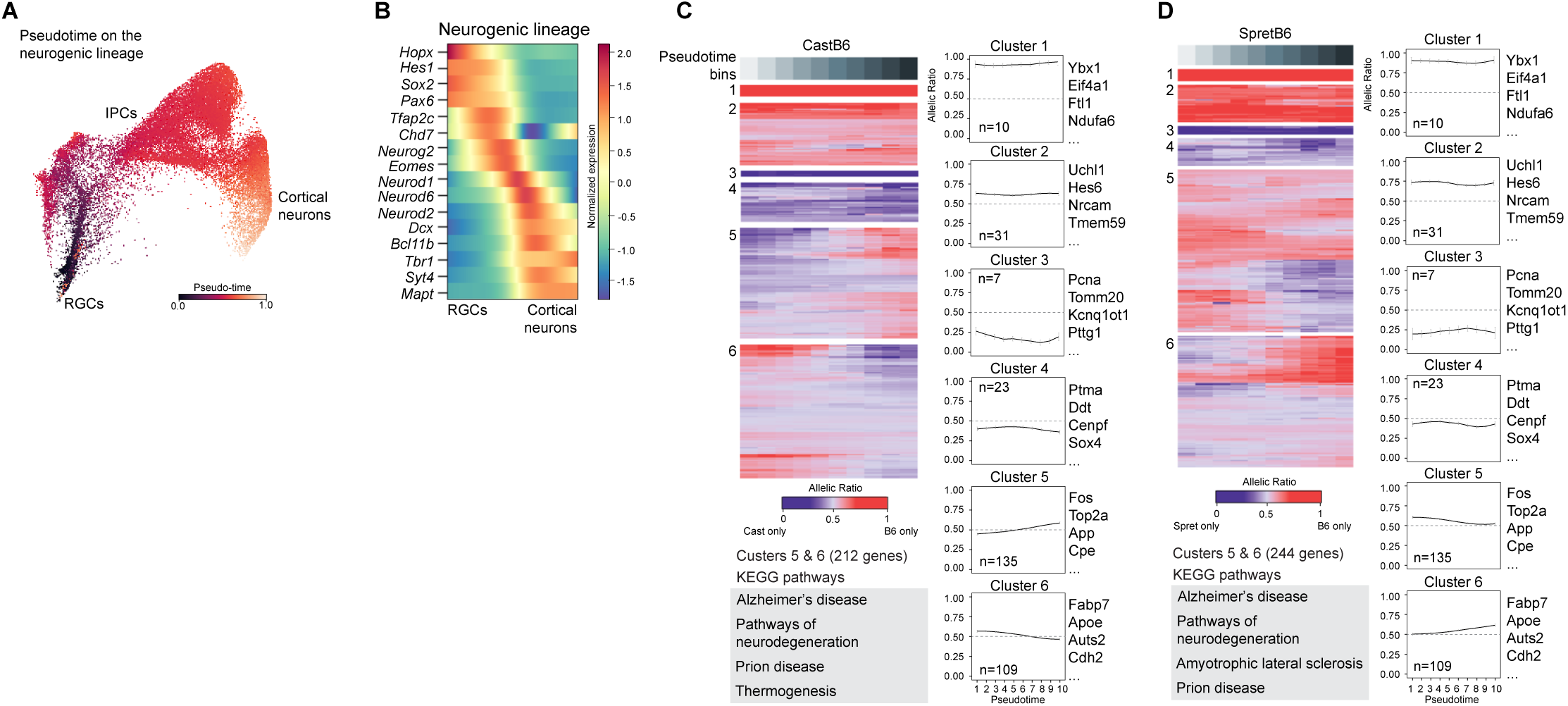
Pseudo-time analyses on cortical organoids inform about allelic imbalances in F1 hybrids along cortical neuron development with single-cell resolution. (**A**) Force-directed layout representation of pseudo-time analyses on the neurogenic lineage as detected by scRNA-seq on day12 organoids using Palantir32. Only the cells identified as RGCs, IPCs, and cortical neurons were kept, and the starting and ending points were manually annotated based on biological knowledge. (**B**) Gene expression dynamics across the neurogenic lineage. Individual cells were ordered based on pseudotime coordinates, from 0 to 1. (**C-D**) Heatmap of allelic ratio trends along the neurogenic pseudotime, for CastB6 (C) and SpretB6 (D) organoids, for 315 genes with significant association (p-value < 0.05, GAM model fit) in at least one background, grouped using hierarchical clustering. Values are averaged within 10 pseudotime bins. Line plots (right) show the average trend. The more significantly enriched pathways in clusters 5 and 6, as detected by KEGG pathway analyses, are displayed under the heatmaps. For extended information on KEGG pathways analyses, see Supplementary Figure 6.

Within this gene set, we identified 58-65% genes that exhibit significant allelic changes along cortical neurogenesis (FDR<0.05). This may be caused by shifts in allelic imbalance and/or variance between different stages of differentiation. Changes in allelic distribution over pseudotime for specific genes can be visualized via a ShinyApp (organoidallele.victorchang.edu.au). To capture common trends of allelic imbalance along differentiation time, we modeled the allelic trajectory of each gene and grouped those genes that were changing in a coordinated manner using hierarchical clustering (see Methods). We identified 315 genes that showed an allelic trend during cortical neuron differentiation in at least one genetic background (p-value < 0.05, GAM model**, Supplementary table 8**). The majority of these genes were identified as dynamic in multiple genetic backgrounds (Clusters 5 and 6 in **Fig. 7C-D** and **Fig. S6A-B**), a subset exhibited consistent allelic biases across all stages of differentiation in one hybrid strain but not another (Clusters 1-4 of **Fig. 7C-D** and **Fig.S6A-B, Supplementary table 9**). As expected, many genes with highly consistently biased allelic expression are imprinted genes (e.g. *Meg3*, *Kcnq1ot1*) or are genes on the X chromosome. Interestingly, the genes with different dynamics of allelic imbalance across pseudotime, include genes known to be important for RGCs and/or neuronal differentiation such as *Nfib*, *Jun*, *Apc*, or *Nrxn1*. Downstream pathway analyses on Clusters 5 and 6 also identified an overrepresentation of genes associated with neurological disorders, in particular Alzheimer’s disease (FDR < 0.05, **Fig. S6C**).

Our findings reveal that allelic expression dynamics are not static but can vary significantly during cortical neuron differentiation, reflecting state-specific regulatory mechanisms. This underscores the complex interplay between genetic background and developmental context in shaping gene expression dynamics during cellular differentiation, providing new insights into how genetic diversity may contribute to intra-population and inter-individual differences in brain development and susceptibility to neurological disorders.

## DISCUSSION

Here, we developed a new protocol to reproducibly generate cerebral cortex organoids from mouse EpiSCs. Mouse cortical organoids recapitulate key features of embryonic and early postnatal mouse cortical development at a tempo similar to *in vivo* development. This will be a valuable new experimental model for studying molecular and cellular mechanisms of cerebral cortex development that can complement *in vivo* studies in mouse as well as human cortical organoid models. Using this protocol, we successfully generated cortical organoids from 18 distinct cell lines from 5 different genetic backgrounds, including F1 hybrids between the common laboratory strain C57BL/6J and 4 wild-derived inbred strains spanning ∼1M years of evolutionary divergence. Using scRNA-seq and a novel computational pipeline for quantification of allelic imbalance (ASPEN), we performed the first comprehensive assessment of the extent of cis-regulatory variation in gene expression across diverse mouse subspecies during cortical neurogenesis with cell-type resolution. This revealed significant cis-regulatory changes between distinct inbred strains, many of which impact gene expression in a cell-type and developmental stage-specific manner. Together, these new experimental methods, F1 hybrid pluripotent stem cell lines, and computational approaches represent a powerful platform for examining molecular and cellular mechanisms of cerebral cortex development and evolution.

Our work builds on pioneering studies that developed the first methods for generating neural organoids from pluripotent stem cells cultured in the naïve pluripotent state^10,16^. Although these studies served as a critical foundation for the subsequent development of human neural organoids, these specific protocols proved difficult to implement and thus have not been widely used. In addition, it was not possible to fully pattern these organoids towards a cortical fate, and thus it was necessary to use a reporter line for cortical cells (*Foxg1* reporter PSCs) to isolate correctly-patterned cells by FACS and re-aggregate these nascent cortical cells to generate cortical organoids^10^. It is now appreciated that this re-aggregation process can potentially have a negative impact on the development of proper tissue architecture and cortical RGC cell fate specification^75^. We recently demonstrated that starting from mouse pluripotent stem cells cultured in the primed pluripotent state (EpiSCs - analogous to the pluripotent state commonly used for human PSC culture) can dramatically improve the reproducibility and robustness of mouse neural organoid generation^18^. In this study, we identified patterning conditions that allow for consistent generation of neural organoids that are nearly homogeneously patterned to a cerebral cortical identity, overcoming critical issues with previous protocols that have limited the impact of mouse cortical organoids as a model system for studying molecular and cellular mechanisms of cortical development. This protocol should thus be of broad interest to neurodevelopmental and stem cell biologists seeking experimentally tractable models of cerebral cortex development.

One of the next frontiers in modeling brain development with organoid systems is to attempt to recapitulate features of early postnatal development, such as neuronal maturation^76^. However, these studies have proven difficult in human organoid models because of the protracted timeline of human cortical development compared to mouse. For example, recent reports indicate that human cortical organoids take at least 250 days to begin to partially recapitulate some features of early postnatal cortical development, roughly similar to the time of human gestation^77^. Mouse cortical organoids can potentially be a valuable complement to studies of organoid maturation, because they develop and mature much more rapidly than human organoids. For example, we see evidence of acquisition of molecular/cellular biomarkers of maturation in mouse cortical organoids in ∼4-6 weeks, including accumulation of CpA methylation in cortical neurons, expression of postnatal gene programs in astrocytes, and differentiation and myelination by cortical oligodendrocytes^49^. In future studies, it will be interesting to characterize the extent of maturation that can be achieved in this model in greater depth to determine how this compares to what is possible with human brain organoids. If more advanced stages of maturation can be achieved, this could enable more accurate modeling of neuronal aging or mechanisms of neurodegenerative disease *in vitro*, which have both proven difficult in human organoid models^76^.

To enable the study of how cis-regulatory variation contributes to phenotypic differences in cerebral cortex development across distinct inbred mouse strains we generated and extensively characterized a new panel of F1 hybrid EpiSCs. Although we focused on characterizing cis-regulatory variation in gene expression in this study, these cell lines represent a powerful experimental model that can be used to study cis-regulatory variation in other developing cell types. Along these lines, we show that these EpiSC lines can also be rapidly and efficiently differentiated into endodermal and mesodermal lineages. Given the difficulty of acquiring, housing, and routinely breeding several of these wild-derived inbred strains, such experiments would be very difficult to perform in developing embryos. The combination of improved directed differentiation protocols, well-characterized cell lines, scalable scRNA-seq and/or multi-omic methods, and new computational tools for allele-specific mapping and quantification (ASPEN) open up exciting new possibilities for mechanistic evolutionary developmental biology studies in genetically diverse inbred and outbred mice^78,79^.

## LIMITATIONS OF THE CURRENT STUDY

Mouse cortical organoids have many of the same limitations as human cortical organoids for modeling cerebral cortex development^80^. For example, mouse cortical organoids generated with our current protocol generate fewer upper-layer cortical neurons than what is observed in the mouse embryo. A similar phenomenon has been observed in human cortical organoid models^81^. Given that cortical neurons with upper-layer identities are mostly generated from RGCs after the generation of deep-layer cells, this could result from premature transition of RGCs to gliogenesis, perhaps due to improper or insufficient environmental cues in the *in vitro* environment. Another possibility is that upper-layer neurons are born properly from RGCs and then undergo apoptosis at a higher rate than *in vivo* due to the lack of appropriate cues to promote their survival, migration, or maturation. A related issue is that cortical organoids lack cells that are derived from other regions of the developing embryo, including GABAergic interneurons, endothelial cells, microglia, and meninges, all of which engage in important intercellular interactions with cortical neurons, astrocytes, and OPCs that facilitate their growth and differentiation^82^. These non-neural cell types would need to be produced separately using a distinct differentiation protocol, and then transplanted back into cortical organoids or aggregated together with cortical organoids to make an assembloid^83^.

We have not performed in depth characterization of cortical organoids cultured for longer than ∼3 weeks. It is likely that further optimization will be necessary to identify optimal conditions for long-term culture of cortical organoids and to potentially promote further maturation of cortical neurons, as has been the case for human organoid models^77,84^. Optimized long-term culture conditions will help evaluate the functional and physiological maturation of neurons in cortical organoids to determine the extent to which they can recapitulate physiological properties of immature neural circuits that develop after birth.

A current limitation of the allele-specific mapping and transcription quantification analyses is the accuracy of the reference genomes available for each of the wild-derived inbred strains. For the analyses in this study, we used the genome builds generated in^55^ (see Methods). Relatively recently, improved genome builds based on long-read sequencing methods have been generated, but they are not yet available for each of the inbred strains used here^85^. These higher-quality genomes will further improve the accuracy of allele-specific mapping and quantification, and can help to reduce mapping artifacts. In addition, we used 10X Genomics 3’ Single Cell platform to generate scRNA-seq/snRNA-seq data in this study, which leads to a 3’ bias in mRNA coverage. In future studies, utilizing a scRNA-seq method that captures transcripts using random hexamers instead of poly-dT sequences could also further improve allelic quantification and would make it possible to more comprehensively assess differential mRNA splicing between alleles^86–88^.

## Supporting information

Key resources table

Supplementary experimental methods

## ACKNOWLEDGEMENTS

We would like to thank all of the members of the Vierbuchen laboratory for helpful discussions and feedback. This study would not have been possible without the core facilities at MSKCC. We would like to specifically thank Murray Tipping and Michael Galiano (Molecular Cytology Core, MSKCC) and members of the Integrated Genomics Operation Core (MSKCC) for their advice and assistance with experiments and data analysis throughout the course of this project. In addition, we would like to thank Dr. Ronan Chaligne and Dr Roshan Sharma (Single Cell Research Initiative, MSKCC) for guidance and advice related to single cell RNA and scATAC-seq data analysis. We would like to thank our MSKCC colleagues Dr. Anna Katerina Hadjantonakis, Dr. Lorenz Studer, Dr. Alexandra Joyner, Dr. Ryan Walsh, for helpful discussions and advice throughout the study. We would like to thank Dr. Laura Santini (MSKCC) and Dr. Nan Yang (Icahn School of Medicine, Mount Sinai) for providing comments on the manuscript.

We would also like to acknowledge Qing Wang and David Humphreys for computational research assistance, National Computational Infrastructure (NCI) and Victor Chang Innovation Centre for support with computation.

## AUTHOR CONTRIBUTIONS

*Conceptualization*: D.M.C., M.T.I., V.P., E.S.W., T.V.

*Methodology*: D.M.C., M.T.I., V.P., E.S.W., T.V.

*Formal Analysis*: D.M.C., V.P., M.G.Y., E.S.W.

*Investigation*: D.M.C., M.T.I., Z.B., S.D.

*Resources*: M.S.

*Data Curation*: D.M.C., V.P., E.S.W.

*Writing – original draft*: D.M.C., V.P., E.S.W., T.V.

*Writing – review and editing*: D.M.C., V.P., E.S.W., T.V.

*Supervision*: E.S.W., T.V.

*Project Administration*: E.S.W., T.V.

*Funding Acquisition*: E.S.W., T.V.

## DATA AVAILABILITY

Bulk RNA-seq data from EpiSCs is deposited with the Gene Expression Omnibus (GEO) GSE268329. Single-cell RNA-seq data from EpiSCs is deposited at GEO, GSE268332. Single-cell RNA-seq data from day 12 cortical organoids is deposited at GEO, GSE268332. Single-nuclei RNA-seq data from day 18 cortical organoids is deposited at GEO, GSE268332. Single-cell ATAC-seq data from day 18 cortical organoids is deposited at GEO, GSE268331.

## MATERIALS AND METHODS

### Summary

For information about antibodies used for immunofluorescence staining and detailed information about reagents, see **KEY RESOURCES TABLE**.

For detailed information and quality control data for each cell line, see **Supplementary Table 1**

### Mouse iPSCs culture and conversion to EpiSCs

All mouse iPSC lines were maintained on gelatin-coated dishes with irradiated mouse embryonic fibroblast (MEFs) feeder cells using serum/LIF media comprised of DMEM (high glucose, GlutaMAX, HEPES), 1% nonessential amino acids, 1% sodium pyruvate, 1% penicillin-streptomycin, 0.1% 2-mercaptoethanol, 10% fetal bovine serum and 1000 units/mL ESGRO LIF. All iPSC lines were cultured in serum/LIF media. For bulk RNA and ATAC-seq, the iPSCs were cultured in feeder-free conditions using serum/LIF media with 2i containing 3 μM CHIR99201 and 1 μM PD0325901. Media was changed daily, and iPSCs were passaged upon 70% confluence at a 1:6 ratio using TrypLE. For iPSC-to-EpiSC conversion, iPSCs were lifted using 0.1 u/μL Collagenase IV, centrifuged, and dissociated into a single-cell solution using Accutase. The iPSC suspension was plated on gelatin-coated dishes with irradiated MEFs at a density of ∼50,000-100,000 cells per cm^2^. EpiSC media consisted of N2B27 supplemented with 20 ng/mL activin A, 12.5 ng/mL heat stable bFGF and Wnt inhibitor (175 nM NVP-TNKS656). EpiSC media was changed daily, and cells were passaged every ∼48 hours at a 1:6/1:12 ratio using 0.1 u/μL collagenase IV followed by dissociation with Accutase into small clumps of 3-5 cells. Both iPSCs and EpiSCs were frequently tested for mycoplasma by PCR (Mycoplasma PCR Detection Kit, G238, abm).

### EpiSCs Karyotype

EpiSC lines were karyotyped at P12P5 (7 passages as ESCs and 5 passages as EpiSCs) (**Supplementary Table 1**). Dissociated EpiSCs were seeded at a dilution to achieve ∼70% confluency at the time of karyotyping. EpiSC cultures were incubated with 0.05 mg/mL KaryoMAXTM ColcemidTM solution in PBS (Thermo Fisher Scientific) for 30-40 minutes at room temperature to arrest the cell cycle in metaphase. EpiSCs were detached and dissociated with 100µL Trypsin-EDTA solution (0.25%) with EDTA (0.02%), neutralized with PBS solution, and centrifuged to form a pellet. The cell pellet was resuspended with 8mL pre-warmed 0.075 M KCl, incubated at 37 °C for 8 minutes, and centrifuged. The cell pellet was fixed with 5 mL methanol:glacial acetic acid (3:1) fixative solution for 5-10 minutes at room temperature, washed two times with the same amount of fixative solution, and centrifuged. Next, the cells were resuspended in 500 mL fixative solution and released as a single drop on a slide vertically with the end about 10 cm above the slide. Slides were air-dried at room temperature, incubated at 37 °C for 3 days, and mounted with VECTASHIELD Antifade Mounting Medium with DAPI. For each sample, chromosome analysis was performed on a minimum of 20 metaphase cells that were fully karyotyped using a Nikon confocal microscope with 100x objective.

### Generation of iPSCs from F1 hybrid mouse embryonic fibroblasts

An optimized iPSC reprogramming protocol developed by Vidal et al, 2014 was used^89^. Briefly, F1 hybrid MEFs were isolated from E13.5 - E14.5 embryos. MEFs were cultured and expanded in MEF media (DMEM with glutamine, 1X penicillin-streptomycin, and 10% fetal calf serum). For reprogramming, MEFs were dissociated into a single-cell suspension using TrypLE and seeded at 5,000 cells/well in MEF media in a 96-well plate coated with 0.1% gelatin. After 24 hours, MEFs were transduced with concentrated, tittered TetO-FUW-OSKM (Addgene, plasmid #20321) and FUW-M2rtTA (Addgene, plasmid #20342) lentiviruses in 100 µL MEF media with 8 µg/mL polybrene. After 8 hours of incubation with lentiviral media, the media was changed to MEF media for the cells to recover for 16 additional hours. Transduced MEFs were then dissociated using TrypLE and replated onto a 10 cm dish with irradiated feeders in MEF media. Reprogramming was initiated by switching the media to mouse serum/LIF ESC media (see iPSCs culture section) containing 1 µg/mL doxycycline and the 3c cocktail (50 ng/µL ascorbic acid, 250 nM iALK5, 3 µM CHIR99021). Media was changed every 2-3 days with fresh 3c/Dox Serum/Lif ESC media. After seven days of 3c/doxycycline treatment, media was removed, cells were washed twice with PBS to ensure complete removal of doxycycline and cultured in Serum/LIF ESC media containing 50 ng/µL ascorbic acid. Individual colonies were picked following a minimum of 72 hours of culture after the removal of doxycycline. Each iPSC clonal cell line was expanded for 2-3 passages, converted to EpiSCs, and the remainder was frozen down in Serum/LIF ESC media with 10% DMSO.

### Definitive endoderm and paraxial mesoderm differentiation

Definitive endoderm differentiation was performed as previously described^90^. Briefly, EpiSCs were detached using Collagenase IV (0.1 u/μL) and then dissociated into a single cell suspension using Accutase. EpiSCs were plated at a density of 110,000 cells/cm^2^ in chemically defined media (CDM) containing 50% IMDM, 50% Ham’s F12 Nutrient Mix with Glutamax, 1% chemically defined lipid concentrate, 450 µM Monothioglycerol, 1% polyvinyl alcohol (w/v), 15 µg/mL Apo-transferrin, 0.5% Glutamax, 0.7 µg/mL Insulin and supplemented with 20 ng/mL Activin A, 12.5 ng/mL heat stable bFGF, 175 nM NVP-TNKS656, 1% knockout serum replacement (KSR), 5 µM Emricasan, and 50 nM Chroman1. Plating media was removed 6 hours after seeding, cells were washed gently with PBS without calcium or magnesium (PBS^-/-^) and the media was changed to CDM supplemented with 3 μM CHIR99201 and 40 ng/mL Activin A. 16 hours after the first media change, cells were washed with PBS^-/-^ and the media was replaced with CDM supplemented with 100 ng/mL Activin A and 100 nM LDN-193189. After 24 hours (46 total hours after cells were initially seeded), cells were fixed for immunostaining.

For paraxial mesoderm differentiation, EpiSCs were plated at the same density and with the same media as for definitive endoderm differentiation. Similarly, plating media was removed 6 hours after seeding, and cells were washed once with PBS^-/-^. The media was changed to CDM supplemented with 3 µM CHIR99201 and 25 ng/mL heat-stable bFGF. The media was replaced after 16 hours. The cells were then fixed for immunostaining after a total of 40 hr in differentiation media.

### Immunostaining of adherent cells

For immunostaining of adherent cells, cells were rinsed twice with PBS^-/-^ and fixed with 4% paraformaldehyde for 30 minutes at room temperature. After fixing, cells were washed twice with PBS^-/-^ and permeabilized with PBS^-/-^ containing 1% Triton-X for 10 minutes at room temperature and then blocked for 30 minutes with PBS^-/-^ containing 0.3% Triton-X and 5% fetal bovine serum (FBS). Primary antibodies were diluted in PBS containing 0.1% Triton-X (PBST) with 1% FBS and incubated overnight at 4 °C. The primary antibodies used to immuno-stain the adherent cells are rabbit anti-SOX2 (Sigma-Millipore, 1:500), rabbit anti-NANOG (Cell Signaling Technologies, 1:500), mouse anti-OCT4 (Santa Cruz, 1:500), goat anti-KLF4 (1:200), goat anti-TBX6 (R&D systems, 1:200), rabbit anti-Brachyury/T (Abcam, 1:1000), goat anti-SOX17 (R&D systems, 1:200), rabbit anti-FOXA2 (Abcam, 1:500). Detailed information about the antibodies can be found in **KEY RESOURCES TABLE**. Following overnight incubation and three 10-minute washes with PBST, cells were incubated with secondary antibodies for 2 hours at room temperature. All secondary antibodies were diluted at 1:500 in PBST with 1% FBS. Cells were rinsed for three 10-minute washes with PBST before DAPI staining (1 μM, diluted in PBST) for 15 minutes at room temperature. Cells were then rinsed with PBST and imaged using a Leica Dmi8.

### Protocol for generating neocortical organoids

For cortical organoid generation, EpiSC colonies were detached using collagenase IV (0.5 u/μL), washed once with PBS^-/-^, and then dissociated into a single-cell suspension using Accutase or TrypLE (Thermo Scientific). EpiSCs were then seeded at 1,000 cells/microwell in an AggreWell^TM^400 plate (34411, STEMCELL Technologies) in embryoid body (EB) formation media containing 50 nM chroman-1, 5 μM emricasan, 400 nM LDN-193189, 10 μM SB-431542, 1 µM cyclopamine and 100 nM LGK-974 in N2B27 (B27 without vitamin A) media. After 24 hours, the EBs were recovered and transferred to the same media but without chroman-1 and emricasan and embedded in Matrigel as previously described (Qian et al., 2018). Briefly, 66.7 μL of EBs and media were mixed with 100 µL of Matrigel using wide-bore tips. The mix was added to a well of a regular 6-well plate without touching the edges of the well and incubated at 37 °C for 30 minutes. After 30 minutes, 3 mL of warm media was added on top. On day 2, the media was changed to plain N2B27 (B27 without vitamin A) with 1 µM cyclopamine and 1% KSR and kept for 24 more hours. On day 3, the media was changed by N2B27 (B27 supplement without vitamin A) containing 1 µM cyclopamine, 3 µM CHIR99201, and 1% KSR. Day 4 organoids were recovered from Matrigel using Cell recovery solution (Corning). Briefly, the wells were washed once with PBS, and 2-3 mL of cell recovery solution was added, while ensuring that the Matrigel dome was efficiently lifted prior to incubation, to ensure efficient organoid recovery. The plates were then incubated for 30 minutes at 4°C. After those 30 minutes, organoids were washed twice in organoid media (N2B27 - B27 supplement without vitamin A - with 20% KSR). Then, the day 4 the cortical organoids were transferred to Petri dishes and kept in N2B27 (B27 supplement without vitamin A) with 20% KSR while shaking at 65 rpm (Infors HT, Celltron benchtop shaker). A step-by-step protocol is provided in the **Supplementary Experimental Methods.**

### Immunofluorescence Staining of organoids

For organoid staining, the organoids were collected in tubes pre-coated with 1% BSA, to avoid attachment. The organoids were washed twice with PBS^-/-^ and fixed with 4% PFA for 45 minutes at 4°C. Afterward, the organoids were washed twice with PBS^-/-^ and permeabilized with PBS containing 0.5% Triton-X for 15 minutes at 4 °C. Then, the organoids were blocked for 15-30 minutes in organoid washing solution (OWS, adapted from^91^), consisting of 0.2% Triton-X, 0.02% SDS, and 0.2% BSA in PBS^-/-^ at 4 °C. The organoids were then incubated in OWS with primary antibodies while shaking overnight at 4 °C. The primary antibodies used to immuno-stain the organoids are rabbit anti-GFP (Abcam, 1:500), rabbit anti-BF1/FOXG1 (Takara, 1:500), rabbit anti-TBR1 (Abcam, 1:500), chicken anti-TBR2 (Millipore, 1:500), rat anti-CTIP2 (Abcam, 1:200), mouse anti-CUX1 (Abcam, 1:200), mouse anti-PAX6 (BD Biosciences, 1:200), mouse anti-SATB2 (Abcam, 1:200), chicken anti-NESTIN (Aves, 1:500), rabbit anti-OLIG2 (Abcam, 1:500), goat anti-POU3F3/BRN1 (Novus Biologicals, 1:100), rat anti-MBP (Novus Biologicals, 1:1000), goat anti-PDGFR*α* (R&D systems, 1:500), rat anti-GFAP (Thermo Fisher Scientific, 1:1000), rabbit anti-S100*β* (Abcam, 1:500), mouse anti-CNP (Atlas Antibodies, 1:1000), chicken anti-NF-H (BioLegend, 1:2000), and rabbit anti-NEUN (Abcam, 1:500). Detailed information about the antibodies can be found in **KEY RESOURCES TABLE**. On the following day, the organoids were washed three times for 30 minutes each with OWS while shaking, and then incubated overnight at 4 °C with secondary antibodies and DAPI (1 μM) in OWS while shaking. All secondary antibodies were diluted at 1:500 in OWS. On the following day, the organoids were once again washed three times with OWS for 1.5 hours while shaking. After that, the organoids were mounted in DeepClear solution (Celexplorer), following the same setup as previously described^90^. The samples were imaged using a confocal Leica SP8, and then analyzed using the Imaris software 10.0.1.

### RNA extraction for RNA-seq

For bulk RNA preparation, about 1 million EpiSCs were detached using collagenase IV (0.5 u/μL), washed once with PBS^-/-^, and then dissociated into a single-cell suspension using Accutase or TrypLE (Thermo Scientific). 500 μL of TRIzol were used to resuspend the pellet and lyse the cells. Phase separation in cells lysed in TRIzol was induced with 50% isopropanol with 0.5% 2-Mercaptoethanol and RNA was extracted from the aqueous phase using the MagMAX mirVana Total RNA Isolation Kit on the KingFisher Flex Magnetic Particle Processor according to the manufacturer’s protocol with 1-2 million cells input. Samples were eluted in 38 μL elution buffer.

### Transcriptome sequencing of EpiSC samples

After RiboGreen quantification and quality control by Agilent BioAnalyzer, 100-500 ng of total RNA with RIN values of 9.5-10 underwent polyA selection and TruSeq library preparation according to instructions provided by Illumina (TruSeq Stranded mRNA LT Kit), with 8 cycles of PCR. Samples were barcoded and run on a NovaSeq 6000 in a PE100 run, using the NovaSeq 6000 S4 Reagent Kit (200 Cycles) (Illumina). An average of 40 million paired reads were generated per sample and the percentage of mRNA bases per sample ranged from 84% to 89%.

### RNA-seq analysis

The resulting fastq files were mapped to transcripts and quantified using salmon^92^ using the mouse genome version mm10. For downstream bulk analyses, Rstudio was used with the packages Deseq2, PCAExplorer and pheatmap^93,94^. Statistical significance was calculated using a Wald test and corrected using the Benjamini-Hochberg method.

### Single-cell RNA sequencing preparation, demultiplexing, and analysis

Multiple organoids from the 4 F1 lines (CastB6, MolfB6, PwkB6 and SpretB6) were collected separately on day 12, washed with PBS^-/-^ and dissociated with Accutase + 10 U/mL DNase I at 37°C until a single-cell suspension was achieved (20-30 minutes). The single-cell suspension was washed several times with PBS and further purified by filtering it through a Flowmi cell strainer (40μm) to remove clumps. Then, the cell number was counted and an equal number of cells from the 4 F1 lines were mixed and used for 10X scRNA-seq (10x Genomics, Single Cell 3’ Kit v3.1, dual index) following the manufacturer’s directions. Combining multiple genetically distinctive lines allowed us to overload the machine (30,000 cells per lane instead of 10,000) since ∼75% of the doublets would be detected as mixes of two different genotypes. A total of 4 clones per F1 line were used, for a total of 16 clones, across 4 different batches.

For scRNA-seq on EpiSCs, the cells were detached using collagenase IV (0.5 u/μL), washed once with PBS^-/-^, and then dissociated into a single-cell suspension using Accutase or TrypLE (Thermo Scientific). After this, the same protocol was applied for organoid or EpiSC scRNA-seq.

We sequenced with enough depth to obtain 40,000 reads per cell, to facilitate allelic mapping. The resulting fastq files were processed with CellRanger (10X Genomics cloud). For demultiplexing, cellsnp-lite and vireo were used^95,96^. The bam file and the barcode file generated by CellRanger were used as input. As VCF file, the VCFs from the reference genomes from the Mouse Genomes Project^56^ - CAST/EiJ, MOLF/EiJ, PWK/PhJ and SPRET/EiJ - were used.

Scanpy was used for downstream analyses^97^. Briefly, only the cells assigned by vireo to one of the four lines were used. Upon downstream clustering, cells within clusters with an abnormally high number of mitochondrial reads or a low number of total reads were excluded. After filtering, ribosomal and Gm-genes were excluded to reduce noise when calling cell-type marker genes.

### Single-nuclei RNA sequencing preparation and analyses

Two different clones of SpretB6 and two biological replicates of B6.129 EpiSCs were used to produce the organoids, which were processed as two different batches on day 18. Single-cell suspension was prepared as for scRNA-seq. After filtering, the cells were kept on ice and we proceeded with nuclear isolation as recommended by 10x Genomics. Briefly, 1 million cells were centrifuged at 300g for 5 minutes at 4 °C. After removal of the supernatant, 100 μL of lysis buffer was added (10 mM Tris-HCl pH 7.4, 10 mM NaCl, 3 mM MgCl_2_, 0.1% Tween-20, 0.1% Nonidet P40 Substitute, 0.01% Digitonin and 1% BSA in nuclease-free water). The cells were resuspended ∼10 times and kept on ice for 5 minutes. Then, 1 mL of washing buffer (10 mM Tris-HCl pH 7.4, 10 mM NaCl, 3 mM MgCl_2,_ 0.1% Tween-20, and 1% BSA in nuclease-free water) was added and the mix was centrifuged at 500g for 5 minutes at 4 °C. Then, the supernatant was discarded, and the nuclei were resuspended in Nuclei Buffer (10x Genomics). 15,000 nuclei were loaded by lane, following the manufacturer’s instructions (10x Genomics, Single Cell 3’ Kit v3.1, dual index), and ∼40,000 reads were sequenced per cell. The resulting datasets were demultiplexed and analyzed following the same guidelines as described for scRNA-seq on day 12 organoids. For comparison of the oligodendrocyte clusters present in day 18 cortical organoids with available datasets, we used the dataset generated by^51^. Briefly, we extracted the gene signature (as detected by bulk RNA-seq) of OPCs in this study after 0, 24, 48, and 72 hours of treatment with Thyroid hormone (T3) (oligodendrocyte progenitor cells, immature oligodendrocytes, mature oligodendrocytes, and myelinating oligodendrocytes, respectively). To compare the astrocyte cells detected by snRNA-seq (d18), we compared their signature with the signature published by Lattke and colleagues^52^. Briefly, we estimated the similarity of their gene signatures (mature VS immature cortical astrocytes), as detected by RNA-seq of *in vivo* astrocytes, with the gene signature of our astrocyte clusters.

### Single nuclei ATAC sequencing preparation and analyses

Two different clones of SpretB6 and two biological replicates of B6.129 EpiSCs were used to produce the organoids, which were processed on two different batches at day 18. The same batches of organoids were used for snRNA-seq and snATAC-seq. The nuclei were isolated as described for snRNA-seq and tagmented as indicated by 10x Genomics (Chromium Next GEM Single Cell ATAC Reagents Kits v2). 10,000 nuclei were loaded per lane and ∼25,000 reads per cell were obtained.

The resulting fastq files were first processed using CellRanger counts following the standard pipeline and the genome version mm10. ArchR^98^ was used for downstream analyses. Only cells with a minimal TSS enrichment score of 10 and more than 10,000 fragments were kept. To reduce the noise caused by the different clones and the influence of the genetic background to call cell types, the samples were corrected per clone using Harmony^99^. The integrated version of macs2 was used to call peaks^100^. getGroupBW was used to generate the bigwig files used in the here-presented tracks, normalizing for reads in the TSS.

### Allele-specific analysis of RNA-seq data (bulk and scRNA-seq)

#### Combined genome generation

The reference C57BL/6J mouse genome (GRCm38, release 68) was concatenated with CAST/EiJ, MOLF/EiJ, PWK/PhJ or SPRET/EiJ genomes to create four combined genomes for read alignment of F1 animals. Strain-specific genomes were generated by incorporating known genetic variants (SNPs and indels) for CAST/EiJ, MOLF/EiJ, PWK/PhJ and SPRET/EiJ genomes, Mouse Genome Project release v5^56^, by substitution into C57BL/6J genome with bcftools consensus (v1.12^101^). Strain-specific gene annotations were created by converting C57BL/6J gene annotation (Ensembl release 102) using UCSC liftOver^102^ with chain files produced by bcftools. Combined genome index files were made using STAR (v2.7.6a^103^).

#### RNA-seq processing

Paired-end fastq files were trimmed using Trimmomatic (v0.39^104^) and then mapped to the combined genome index using STAR (v2.7.6a) allowing for full-length read alignment, without insertions or soft-end clipping, splicing mode was not used (--outBAMcompression 6 -- outFilterMultimapNmax 1 --outFilterMatchNmin 30 --alignIntronMax 1 --alignEndsType EndToEnd -- outSAMattributes NH HI NM MD AS nM --scoreDelOpen -1000 --scoreInsOpen -1000). Reads with any mate mismatches (flagged as NM) and reads overlapping strain-specific diverse regions that are enriched for recently transposed long interspersed nuclear elements (LINEs) and long-terminal repeat (LTR) elements as described by Lilue et al were removed^105^. Reads were counted using Rsubread (v2.4.3^106^) with parameters isPairedEnd=T, allowMultiOverlap=T, countMultiMappingReads=F. To normalise for sequencing depth, size factors were estimated using combined allelic counts with DESeq2 (v1.30.1^93^) and used to adjust allelic counts.

#### scRNA-seq processing

To minimize batch effects, cells from four F1 hybrids were pooled. To separate the strains, we first aligned reads to C57Bl6/J mouse genome (refdata-gex-mm10-2020-A) using Cellranger count (v6.0.0^107^) (--expect-cells=10000 --include-introns). SNP calling was performed in cellsnp-lite (v1.2.3^96^) using strain-specific vcf files. Vireo (v0.5.8^95^) was used to demultiplex cell barcodes for the four strains. Using these strain-resolved cell barcodes, we mapped each set of reads to their respective combined genome index using STARSolo (v2.7.6a^108^) (-- soloType CB_UMI_Simple --soloUMIlen 12 --soloCBstart 1 --soloCBlen 16 --soloUMIstart 17 -- soloCBwhitelist <(zcat ${dir}/align/3M-february-2018.txt.gz) --soloFeatures Gene –outSAMattributes NH HI nM AS CR UR CB UB GX GN sS sQ sM NM MD --outBAMcompression 6 -- outFilterMultimapNmax 1 --outFilterMatchNmin 30 --alignIntronMin 20 --alignIntronMax 20000 -- alignEndsType EndToEnd --outFilterMismatchNoverReadLmax 0.001 --scoreDelOpen -1000 -- scoreInsOpen -1000).

#### Statistical test for allelic imbalance testing

To identify genes with significant allelic imbalance, we assumed the probability of observing allelic ratio follows a beta-binomial distribution,

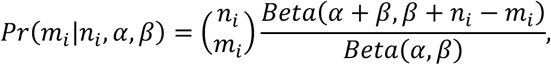

where *m_i_* and
*n_i_* denote reference allele and total allelic counts respectively and parameters α and β guide the shape of the distribution curve. Beta-binomial mean is 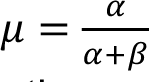 and when α = β, μ = 0.5. Based on that, we consider the following null and alternative hypotheses for absence (*H*_0_) or presence (*H*_1_) of allelic imbalance: *H*_0_: α = β and *H*_1_: α ≠ β. Since we observed a reference allele mapping bias in our data, the initial hypotheses testing procedure required adjustment. We introduced a factor τ = β_*glob*_/α_*glob*_, where α_*glob*_ and β_*glob*_ were estimated by maximum likelihood with optim function in R (R version 4.3.1; R Core Team, 2023) using all genes in dataset, excluding the genes located on sex chromosomes and imprinted genes, to evaluate a general skew towards the reference allele. Then, the test for allelic imbalance is implemented as *H*_0_: α = α × τ and *H*_1_: α ≠ β. Additionally, we implemented a dispersion shrinkage procedure. The motivation for the dispersion adjustment is to improve our ability to detect allelically imbalanced genes in lowly expressed genes and reduce the number of false positive calls among highly expressed genes. For this, we estimate the level of dispersion for each gene as θ̂_*i*_ = 1/(α + β), which is then modelled as a function of mean counts across alleles using locfit function in R as *log*(Dθ̂_*i*_ + 1) = *log*(*n̅_i_*) and predicted values θ̅_*i*_ are obtained. For shrinkage, we used a weighted likelihood approach which was previously used for the Negative Binomial distribution^109–111^. Each gene *i* weighted log-likelihood, ℓ_*WL*_(θ_*i*_), is a combination the individual gene’s log likelihood and a weighted shared log-likelihood

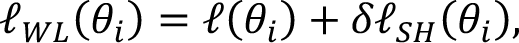

where weight δ is a tuning parameter. We can interpret the weighted likelihood in empirical Bayes terms, with the shared log-likelihood as the prior distribution for θ_*i*_ and the weighted log-likelihood as posterior. Using predicted θ̅_*i*_ values from locfit model as the dispersion priors, the shrunk dispersion is calculated as

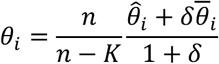

where δ is a tuning parameter, *n* is the sample size and *K* is the number of groups compared. Shrunk dispersion θ_*i*_ is used to correct α and β parameters

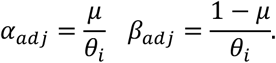

The null and alternative hypotheses to test for allelic imbalance are updated with corrected α and β: *H*_0_: α_*adj*_ = α_*adj*_ × τ and *H*_1_: α_*adj*_ ≠ β_*adj*_. p-values are obtained by a likelihood ratio test and adjusted for the false discovery rate (fdr). For scRNA data, gene counts were pseudo-bulked by clone across the identified cell types, and the allelic imbalance test was performed for each cell type separately.

### Mixed linear models to test for strain and cell type effects in F1 organoids

We modeled the expression of each gene using a generalized linear mixed model with a beta-binomial error distribution. The model included an intercept term, fixed effects for strain and cell type, a fixed interaction effect between strain and cell type, and a random effect for each clone, which is assumed to be Gaussian. We included total gene expression term to all models to account for differences in expression levels. We restricted the values the fixed-effects parameters can take and assessed how well the different models described the different scenarios. We imposed constraints on the range of values that the fixed-effect parameters could take and evaluated the goodness of fit of the models under different scenarios. The models were fitted by maximizing the likelihood, with the Laplace approximation used to integrate out the random effect. The optimization of the marginal likelihood yielded the maximum likelihood estimates of the fixed effects and variance components. This was done using ‘glmmTMB’ v.1.1.9 (https://github.com/glmmTMB/glmmTMB) in R.

To determine if a gene exhibited differential allelic expression between strains, we evaluated the fit of the data to two competing models. Only genes with all four models converged were used for this analysis. The null model constrained the strain term and the interaction term between strain and cell type to be zero. In contrast, the alternative model allowed these terms to be freely estimated from the data. Bayesian Information Criterion (BIC) was used to compare the models’ relative fit to the data. A BIC difference above ten was used to determine the model that best fits the gene. Similarly, to test for differences due to cell types, we set the cell type terms to zero and allowed the estimation of other model components.

### Statistical test for allelic changes during differentiation

To detect genes that undergo allelic expression changes across cortical neurogenesis, we assume that the probability of allelic ratio follows a mixture of beta-binomial distributions. If allelic ratio is maintained across cell types (as captured along the pseudotime vector), α and β parameters of beta-binomial distribution, measured across different pseudotime points, would remain the same, and if alleles are used at different rates, the estimated parameters would vary between the groups. We formulate the test as 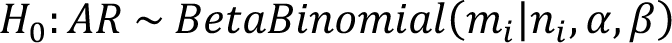 and 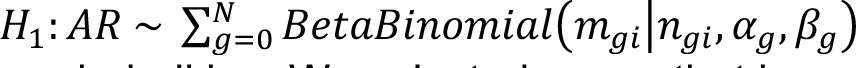. The test was performed on the scRNA data without pseudo-bulking. We selected genes that have at least 5 UMI counts across at least 15 cells. The counts were divided into equal-sized bins across pseudotime trajectory and α and β estimates were obtained across all observations and within each bin. Significance was determined by a likelihood ratio test and adjusted for the false discovery rate.

### Identifying allelic ratio trends across pseudotime

To model allelic ratio trend along pseudotime, we selected genes with a minimum of 5 reads in at least 15 cells from the neurogenic lineage (RGCs, IPCs, and cortical neurons). Pseudotime vector was divided into ten equally sized bins, and the mean allelic ratio was calculated per bin for each gene. Aggregated allelic ratios were modeled as a function of pseudotime with generalized additive model (GAM) using gamlss (v5.4-14) package^112^. For genes with evidence of association between allelic ratio and pseudotime (p-value < 0.05) in at least one strain, we obtained the predicted values and imputed those missing in some pseudotime bins using missForest (v1.5)^113^. Clustering was performed using unsupervised hierarchical clustering with Ward linkage and Euclidean distance metric using hclust in R (R version 4.3.1; R Core Team, 2023).

### Functional term enrichment analyses

Gene set enrichment analyses were performed with ClusterProfiler (v4.8.3^114^) using the org.Mm.eg.db database (v3.17.0^115^). After allelic imbalance test for each cell type, genes were ranked by the absolute -log10 transformed fdr values. These pre-ranked gene lists were analysed using the gseGO function (ont =“ALL”, by = “fgsea”). GO overlap and KEGG pathway enrichment analyses were performed on a subset of genes with changing allelic biases across pseudotime relative to a background set of all expressed genes, using the enrichGO and enrichKEGG functions respectively.

## ADDITIONAL SUPPLEMENTARY MATERIAL

**Supplementary Experimental Methods.** Step-by-step protocol for generation of cortical organoids

**Supplementary Table 1.** Description of all the cell lines used in this study.

**Supplementary Table 2.** Allelic imbalance test results bulk and pseudobulk comparison EpiSCs.

**Supplementary Table 3.** Allelic measurements and related statistics in day 12 CastB6 cortical organoids, subdivided by cell type.

**Supplementary Table 4.** Allelic measurements and related statistics in day 12 MolfB6 cortical organoids, subdivided by cell type.

**Supplementary Table 5** Allelic measurements and related statistics in day 12 PwkB6 cortical organoids, subdivided by cell type.

**Supplementary Table 6.** Allelic measurements and related statistics in day 12 SpretB6 cortical organoids, subdivided by cell type.

**Supplementary Table 7.** BIC model testing.

**Supplementary Table 8.** Related to Figure 7 and Supplementary Figure 6, GAM model.

**Supplementary Table 9.** Related to Figure 7 and Supplementary Figure 6, genes within each cluster.

## REFERENCES

1. Hodge, R. D. et al. Conserved cell types with divergent features in human versus mouse cortex. Nature 573, 61–68 (2019).

2. Miller, D. J., Bhaduri, A., Sestan, N. & Kriegstein, A. Shared and derived features of cellular diversity in the human cerebral cortex. Curr Opin Neurobiol 56, 117–124 (2019).

3. Cadwell, C. R., Bhaduri, A., Mostajo-Radji, M. A., Keefe, M. G. & Nowakowski, T. J. Development and Arealization of the Cerebral Cortex. Neuron 103, 980–1004 (2019).

4. Greig, L. C., Woodworth, M. B., Galazo, M. J., Padmanabhan, H. & Macklis, J. D. Molecular logic of neocortical projection neuron specification, development and diversity. Nat Rev Neurosci 14, 755–769 (2013).

5. Di Bella, D. J., Domínguez-Iturza, N., Brown, J. R. & Arlotta, P. Making Ramón y Cajal proud: Development of cell identity and diversity in the cerebral cortex. Neuron 112, 2091–2111 (2024).

6. Huang, Z. J. Toward a genetic dissection of cortical circuits in the mouse. Neuron 83, 1284– 1302 (2014).

7. Fair, T. & Pollen, A. A. Genetic architecture of human brain evolution. Curr Opin Neurobiol 80, 102710 (2023).

8. Molnár, Z. et al. Evolution and development of the mammalian cerebral cortex. Brain Behav Evol 83, 126–139 (2014).

9. Kadoshima, T., et al. Self-organization of axial polarity, inside-out layer pattern, and species-specific progenitor dynamics in human ES cell–derived neocortex. Proc. Natl. Acad. Sci. U.S.A. 110, 20284–20289 (2013).

10. Eiraku, M. et al. Self-Organized Formation of Polarized Cortical Tissues from ESCs and Its Active Manipulation by Extrinsic Signals. Cell Stem Cell 3, 519–532 (2008).

11. Lancaster, M. A. et al. Cerebral organoids model human brain development and microcephaly. Nature 501, 373–379 (2013).

12. Di Lullo, E. & Kriegstein, A. R. The use of brain organoids to investigate neural development and disease. Nat Rev Neurosci 18, 573–584 (2017).

13. Kelley, K. W. & Pașca, S. P. Human brain organogenesis: Toward a cellular understanding of development and disease. Cell 185, 42–61 (2022).

14. Corsini, N. S. & Knoblich, J. A. Human organoids: New strategies and methods for analyzing human development and disease. Cell 185, 2756–2769 (2022).

15. Chiaradia, I. & Lancaster, M. A. Brain organoids for the study of human neurobiology at the interface of in vitro and in vivo. Nat Neurosci 23, 1496–1508 (2020).

16. Nasu, M. et al. Robust formation and maintenance of continuous stratified cortical neuroepithelium by laminin-containing matrix in mouse ES cell culture. PLoS One 7, e53024 (2012).

17. Watanabe, K. et al. Directed differentiation of telencephalic precursors from embryonic stem cells. Nat Neurosci 8, 288–296 (2005).

18. Medina-Cano, D. et al. Rapid and robust directed differentiation of mouse epiblast stem cells into definitive endoderm and forebrain organoids. Development 149, dev200561 (2022).

19. Li, C. et al. Single-cell brain organoid screening identifies developmental defects in autism. Nature 621, 373–380 (2023).

20. Meng, X. et al. Assembloid CRISPR screens reveal impact of disease genes in human neurodevelopment. Nature 622, 359–366 (2023).

21. Fleck, J. S. et al. Inferring and perturbing cell fate regulomes in human brain organoids. Nature 621, 365–372 (2023).

22. Birey, F. et al. Dissecting the molecular basis of human interneuron migration in forebrain assembloids from Timothy syndrome. Cell Stem Cell 29, 248–264.e7 (2022).

23. Swanzey, E., O’Connor, C. & Reinholdt, L. G. Mouse Genetic Reference Populations: Cellular Platforms for Integrative Systems Genetics. Trends Genet 37, 251–265 (2021).

24. Groza, T. et al. The International Mouse Phenotyping Consortium: comprehensive knockout phenotyping underpinning the study of human disease. Nucleic Acids Res 51, D1038–D1045 (2023).

25. Saul, M. C., Philip, V. M., Reinholdt, L. G., Center for Systems Neurogenetics of Addiction & Chesler, E. J. High-Diversity Mouse Populations for Complex Traits. Trends Genet 35, 501–514 (2019).

26. Wang, N. et al. Variability and heritability of mouse brain structure: Microscopic MRI atlases and connectomes for diverse strains. Neuroimage 222, 117274 (2020).

27. Morris, J. A. et al. Divergent and nonuniform gene expression patterns in mouse brain. Proc Natl Acad Sci U S A 107, 19049–19054 (2010).

28. Wahlsten, D., Bachmanov, A., Finn, D. A. & Crabbe, J. C. Stability of inbred mouse strain differences in behavior and brain size between laboratories and across decades. Proc Natl Acad Sci U S A 103, 16364–16369 (2006).

29. Paşca, A. M. et al. Functional cortical neurons and astrocytes from human pluripotent stem cells in 3D culture. Nat Methods 12, 671–678 (2015).

30. Qian, X. et al. Brain-Region-Specific Organoids Using Mini-bioreactors for Modeling ZIKV Exposure. Cell 165, 1238–1254 (2016).

31. Kiecker, C. & Lumsden, A. The role of organizers in patterning the nervous system. Annu Rev Neurosci 35, 347–367 (2012).

32. Hébert, J. M. & Fishell, G. The genetics of early telencephalon patterning: some assembly required. Nat Rev Neurosci 9, 678–685 (2008).

33. Rifes, P. et al. Modeling neural tube development by differentiation of human embryonic stem cells in a microfluidic WNT gradient. Nat Biotechnol 38, 1265–1273 (2020).

34. Metzis, V. et al. Nervous System Regionalization Entails Axial Allocation before Neural Differentiation. Cell 175, 1105–1118.e17 (2018).

35. Gaspard, N. et al. An intrinsic mechanism of corticogenesis from embryonic stem cells. Nature 455, 351–357 (2008).

36. Qian, X. et al. Generation of human brain region–specific organoids using a miniaturized spinning bioreactor. Nat Protoc 13, 565–580 (2018).

37. Lancaster, M. A. et al. Guided self-organization and cortical plate formation in human brain organoids. Nat Biotechnol 35, 659–666 (2017).

38. Oeschger, F. M. et al. Gene expression analysis of the embryonic subplate. Cereb Cortex 22, 1343–1359 (2012).

39. Knowles, R., Dehorter, N. & Ellender, T. From Progenitors to Progeny: Shaping Striatal Circuit Development and Function. J Neurosci 41, 9483–9502 (2021).

40. Redmond, S. A. et al. Development of Ependymal and Postnatal Neural Stem Cells and Their Origin from a Common Embryonic Progenitor. Cell Rep 27, 429–441.e3 (2019).

41. Spassky, N. et al. Adult ependymal cells are postmitotic and are derived from radial glial cells during embryogenesis. J Neurosci 25, 10–18 (2005).

42. Hevner, R. F. et al. Tbr1 regulates differentiation of the preplate and layer 6. Neuron 29, 353– 366 (2001).

43. Hoerder-Suabedissen, A. & Molnár, Z. Development, evolution and pathology of neocortical subplate neurons. Nat Rev Neurosci 16, 133–146 (2015).

44. Di Bella, D. J. et al. Molecular logic of cellular diversification in the mouse cerebral cortex. Nature 595, 554–559 (2021).

45. Iwata, R. et al. Mitochondria metabolism sets the species-specific tempo of neuronal development. Science 379, eabn4705 (2023).

46. Ciceri, G. et al. An epigenetic barrier sets the timing of human neuronal maturation. Nature 626, 881–890 (2024).

47. Wallace, J. L. & Pollen, A. A. Human neuronal maturation comes of age: cellular mechanisms and species differences. Nat Rev Neurosci 25, 7–29 (2024).

48. Clemens, A. W. & Gabel, H. W. Emerging Insights into the Distinctive Neuronal Methylome. Trends Genet 36, 816–832 (2020).

49. Lister, R. et al. Global epigenomic reconfiguration during mammalian brain development. Science 341, 1237905 (2013).

50. Shen, Z. et al. Distinct progenitor behavior underlying neocortical gliogenesis related to tumorigenesis. Cell Rep 34, 108853 (2021).

51. Lager, A. M. et al. Rapid functional genetics of the oligodendrocyte lineage using pluripotent stem cells. Nat Commun 9, 3708 (2018).

52. Lattke, M. et al. Extensive transcriptional and chromatin changes underlie astrocyte maturation in vivo and in culture. Nat Commun 12, 4335 (2021).

53. Phifer-Rixey, M. & Nachman, M. W. Insights into mammalian biology from the wild house mouse Mus musculus. Elife 4, e05959 (2015).

54. Yang, M. G., Ling, E., Cowley, C. J., Greenberg, M. E. & Vierbuchen, T. Characterization of sequence determinants of enhancer function using natural genetic variation. Elife 11, e76500 (2022).

55. Lilue, J. et al. Sixteen diverse laboratory mouse reference genomes define strain-specific haplotypes and novel functional loci. Nat Genet 50, 1574–1583 (2018).

56. Keane, T. M. et al. Mouse genomic variation and its effect on phenotypes and gene regulation. Nature 477, 289–294 (2011).

57. Williams, R. W., Strom, R. C., Rice, D. S. & Goldowitz, D. Genetic and environmental control of variation in retinal ganglion cell number in mice. J Neurosci 16, 7193–7205 (1996).

58. Seecharan, D. J., Kulkarni, A. L., Lu, L., Rosen, G. D. & Williams, R. W. Genetic control of interconnected neuronal populations in the mouse primary visual system. J Neurosci 23, 11178– 11188 (2003).

59. Onos, K. D. et al. Enhancing face validity of mouse models of Alzheimer’s disease with natural genetic variation. PLoS Genet 15, e1008155 (2019).

60. Hsiao, K. et al. A Thalamic Orphan Receptor Drives Variability in Short-Term Memory. Cell 183, 522–536.e19 (2020).

61. Wahlsten, D., Metten, P. & Crabbe, J. C. A rating scale for wildness and ease of handling laboratory mice: results for 21 inbred strains tested in two laboratories. Genes Brain Behav 2, 71–79 (2003).

62. Lazzarano, S. et al. Genetic mapping of species differences via in vitro crosses in mouse embryonic stem cells. Proc Natl Acad Sci U S A 115, 3680–3685 (2018).

63. Vidal, S. E., Amlani, B., Chen, T., Tsirigos, A. & Stadtfeld, M. Combinatorial modulation of signaling pathways reveals cell-type-specific requirements for highly efficient and synchronous iPSC reprogramming. Stem Cell Reports 3, 574–584 (2014).

64. Stevenson, K. R., Coolon, J. D. & Wittkopp, P. J. Sources of bias in measures of allele-specific expression derived from RNA-sequence data aligned to a single reference genome. BMC Genomics 14, 536 (2013).

65. Qi, G. & Battle, A. Computational methods for allele-specific expression in single cells. Trends Genet S0168-9525(24)00169–0 (2024) doi:10.1016/j.tig.2024.07.003.

66. Yazar, S. et al. Single-cell eQTL mapping identifies cell type-specific genetic control of autoimmune disease. Science 376, eabf3041 (2022).

67. 1000 Genomes Project Consortium et al. A global reference for human genetic variation. Nature 526, 68–74 (2015).

68. Link, V. M. et al. Analysis of Genetically Diverse Macrophages Reveals Local and Domain-wide Mechanisms that Control Transcription Factor Binding and Function. Cell 173, 1796–1809.e17 (2018).

69. Collins, S. C. et al. Large-scale neuroanatomical study uncovers 198 gene associations in mouse brain morphogenesis. Nat Commun 10, 3465 (2019).

70. Tabbaa, M., Knoll, A. & Levitt, P. Mouse population genetics phenocopies heterogeneity of human Chd8 haploinsufficiency. Neuron 111, 539–556.e5 (2023).

71. Sittig, L. J. et al. Genetic Background Limits Generalizability of Genotype-Phenotype Relationships. Neuron 91, 1253–1259 (2016).

72. Umans, B. D., Battle, A. & Gilad, Y. Where Are the Disease-Associated eQTLs? Trends Genet 37, 109–124 (2021).

73. Long, H. K., Prescott, S. L. & Wysocka, J. Ever-Changing Landscapes: Transcriptional Enhancers in Development and Evolution. Cell 167, 1170–1187 (2016).

74. Setty, M. et al. Characterization of cell fate probabilities in single-cell data with Palantir. Nat Biotechnol 37, 451–460 (2019).

75. Chiaradia, I. et al. Tissue morphology influences the temporal program of human brain organoid development. Cell Stem Cell 30, 1351–1367.e10 (2023).

76. Ciceri, G. & Studer, L. Epigenetic control and manipulation of neuronal maturation timing. Curr Opin Genet Dev 85, 102164 (2024).

77. Gordon, A. et al. Long-term maturation of human cortical organoids matches key early postnatal transitions. Nat Neurosci 24, 331–342 (2021).

78. Skelly, D. A. et al. Mapping the Effects of Genetic Variation on Chromatin State and Gene Expression Reveals Loci That Control Ground State Pluripotency. Cell Stem Cell 27, 459–469.e8 (2020).

79. Glenn, R. A. et al. A PLURIPOTENT STEM CELL PLATFORM FOR IN VITRO SYSTEMS GENETICS STUDIES OF MOUSE DEVELOPMENT. *bioRxiv* 2024.06.06.597758 (2024) doi:10.1101/2024.06.06.597758.

80. Sandoval, S. O. et al. Rigor and reproducibility in human brain organoid research: Where we are and where we need to go. Stem Cell Reports 19, 796–816 (2024).

81. Qian, X., Song, H. & Ming, G.-L. Brain organoids: advances, applications and challenges. Development 146, dev166074 (2019).

82. Stoufflet, J., Tielens, S. & Nguyen, L. Shaping the cerebral cortex by cellular crosstalk. Cell 186, 2733–2747 (2023).

83. Kanton, S. & Paşca, S. P. Human assembloids. Development 149, dev201120 (2022).

84. Giandomenico, S. L., Sutcliffe, M. & Lancaster, M. A. Generation and long-term culture of advanced cerebral organoids for studying later stages of neural development. Nat Protoc 16, 579–602 (2021).

85. Ferraj, A. et al. Resolution of structural variation in diverse mouse genomes reveals chromatin remodeling due to transposable elements. Cell Genom 3, 100291 (2023).

86. Larsson, A. J. M. et al. Genomic encoding of transcriptional burst kinetics. Nature 565, 251–254 (2019).

87. Hagemann-Jensen, M., Ziegenhain, C. & Sandberg, R. Scalable single-cell RNA sequencing from full transcripts with Smart-seq3xpress. Nat Biotechnol 40, 1452–1457 (2022).

88. Rebboah, E. et al. The ENCODE mouse postnatal developmental time course identifies regulatory programs of cell types and cell states. *bioRxiv* 2024.06.12.598567 (2024) doi:10.1101/2024.06.12.598567.

89. Vidal, S. E., Amlani, B., Chen, T., Tsirigos, A. & Stadtfeld, M. Combinatorial Modulation of Signaling Pathways Reveals Cell-Type-Specific Requirements for Highly Efficient and Synchronous iPSC Reprogramming. Stem Cell Reports 3, 574–584 (2014).

90. Medina-Cano, D. et al. Rapid and robust directed differentiation of mouse epiblast stem cells into definitive endoderm and forebrain organoids. *bioRxiv* 2021.12.07.471652 (2021) doi:10.1101/2021.12.07.471652.

91. Dekkers, J. F. et al. High-resolution 3D imaging of fixed and cleared organoids. Nat Protoc 14, 1756–1771 (2019).

92. Patro, R., Duggal, G., Love, M. I., Irizarry, R. A. & Kingsford, C. Salmon provides fast and bias-aware quantification of transcript expression. Nat Methods 14, 417–419 (2017).

93. Love, M. I., Huber, W. & Anders, S. Moderated estimation of fold change and dispersion for RNA-seq data with DESeq2. Genome Biol 15, 550 (2014).

94. Marini, F. & Binder, H. pcaExplorer: an R/Bioconductor package for interacting with RNA-seq principal components. BMC Bioinformatics 20, 331 (2019).

95. Huang, Y., McCarthy, D. J. & Stegle, O. Vireo: Bayesian demultiplexing of pooled single-cell RNA-seq data without genotype reference. Genome Biol 20, 273 (2019).

96. Huang, X. & Huang, Y. Cellsnp-lite: an efficient tool for genotyping single cells. Bioinformatics 37, 4569–4571 (2021).

97. Wolf, F. A., Angerer, P. & Theis, F. J. SCANPY: large-scale single-cell gene expression data analysis. Genome Biol 19, 15 (2018).

98. Granja, J. M. et al. ArchR is a scalable software package for integrative single-cell chromatin accessibility analysis. Nat Genet 53, 403–411 (2021).

99. Korsunsky, I. et al. Fast, sensitive and accurate integration of single-cell data with Harmony. Nat Methods 16, 1289–1296 (2019).

100. Zhang, Y. et al. Model-based Analysis of ChIP-Seq (MACS). Genome Biol 9, R137 (2008).

101. Danecek, P. et al. Twelve years of SAMtools and BCFtools. Gigascience 10, giab008 (2021).

102. Hinrichs, A. S. et al. The UCSC Genome Browser Database: update 2006. Nucleic Acids Res 34, D590–598 (2006).

103. Dobin, A. et al. STAR: ultrafast universal RNA-seq aligner. Bioinformatics 29, 15–21 (2013).

104. Bolger, A. M., Lohse, M. & Usadel, B. Trimmomatic: a flexible trimmer for Illumina sequence data. Bioinformatics 30, 2114–2120 (2014).

105. Lilue, J., Shivalikanjli, A., Adams, D. J. & Keane, T. M. Mouse protein coding diversity: What’s left to discover? PLoS Genet 15, e1008446 (2019).

106. Liao, Y., Smyth, G. K. & Shi, W. The R package Rsubread is easier, faster, cheaper and better for alignment and quantification of RNA sequencing reads. Nucleic Acids Res 47, e47 (2019).

107. Zheng, G. X. Y. et al. Massively parallel digital transcriptional profiling of single cells. Nat Commun 8, 14049 (2017).

108. Kaminow, B., Yunusov, D. & Dobin, A. STARsolo: accurate, fast and versatile mapping/quantification of single-cell and single-nucleus RNA-seq data. 2021.05.05.442755 Preprint at 10.1101/2021.05.05.442755 (2021).

109. Robinson, M. D. & Smyth, G. K. Moderated statistical tests for assessing differences in tag abundance. Bioinformatics 23, 2881–2887 (2007).

110. McCarthy, D. J., Chen, Y. & Smyth, G. K. Differential expression analysis of multifactor RNA-Seq experiments with respect to biological variation. Nucleic Acids Res 40, 4288–4297 (2012).

111. Ruddy, S., Johnson, M. & Purdom, E. Shrinkage of dispersion parameters in the binomial family, with application to differential exon skipping. The Annals of Applied Statistics 10, 690– 725 (2016).

112. Rigby, R. A. & Stasinopoulos, D. M. Generalized Additive Models for Location, Scale and Shape. Journal of the Royal Statistical Society Series C: Applied Statistics 54, 507–554 (2005).

113. Stekhoven, D. J. & Bühlmann, P. MissForest—non-parametric missing value imputation for mixed-type data. Bioinformatics 28, 112–118 (2012).

114. Wu, T. et al. clusterProfiler 4.0: A universal enrichment tool for interpreting omics data. Innovation (Camb) 2, 100141 (2021).

115. Carlson, M. org.Mm.eg.db. Bioconductor 10.18129/B9.BIOC.ORG.MM.EG.DB (2017).

