## Supplementary figures for "A MOUSE ORGANOID PLATFORM FOR MODELING CEREBRAL CORTEX DEVELOPMENT AND CIS-REGULATORY EVOLUTION IN VITRO"

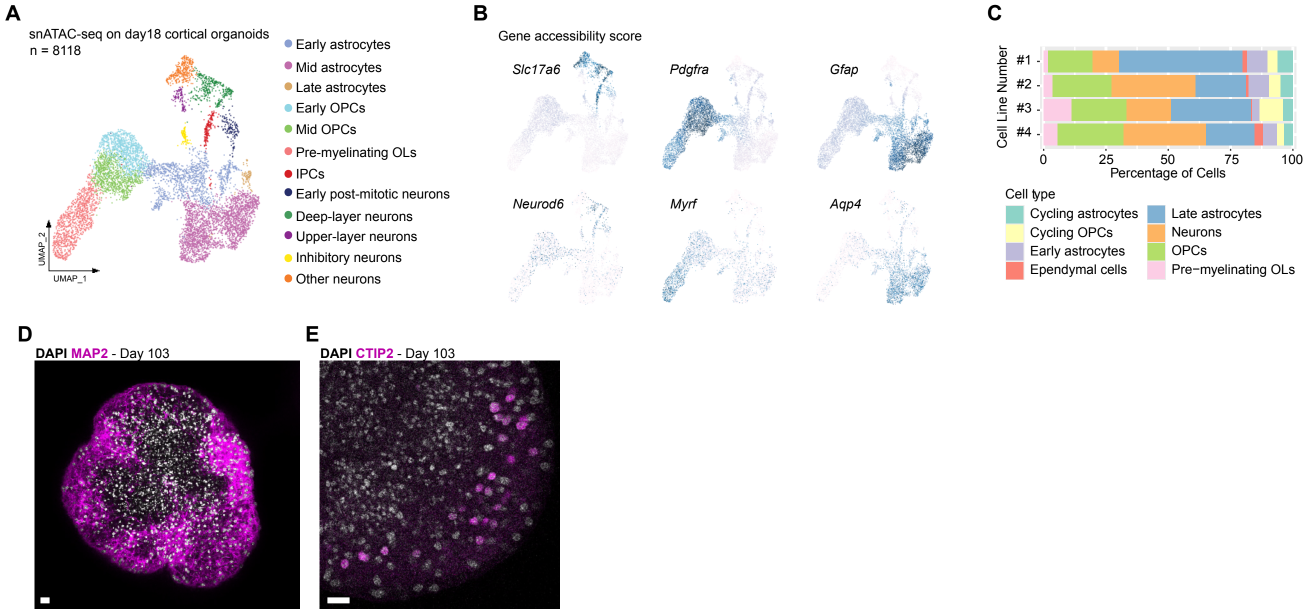


**Supplementary figure 1.** Related to main figure 1. (**A**) Uniform Manifold Approximation and Projection (UMAP) representation of snATAC-seq from d18 cortical organoids, and (**B**) gene accessibility scores for markers of neurons (Slc17a6, Neurod6), oligodendrocytes (Pdgfra, Myrf), and astrocytes (Gfap, Aqp4). (**C**) Contribution of each cell type to d18 organoids across four different cell lines, based on cell type clusters defined from snRNA-seq on d18 cortical organoids. (**D-E**) Immunostaining of d103 organoids for the neuronal markers MAP2 and BCL11B/CTIP2. Scale bars = 20µm. Abbreviations: OPC = Oligodendrocyte progenitor cell, OL = Oligodendrocyte, IPC = intermediate progenitor cell.

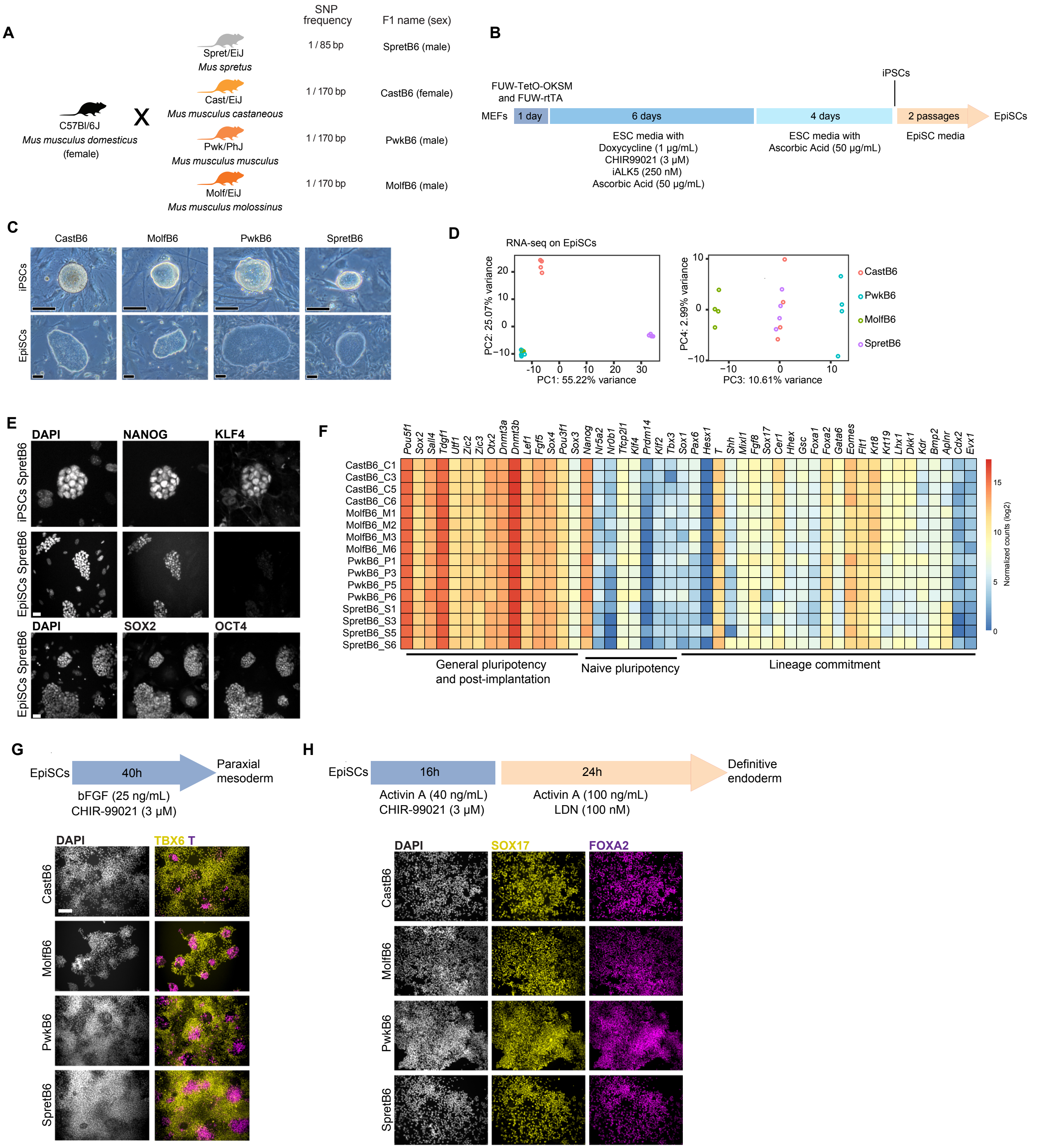


**Supplementary figure 2.** Related to main figure 5. (**A**) Breeding scheme to generate the four F1-hybrid lines used in this study. (**B**) Scheme of the protocol used to generate induced pluripotent stem cells (iPSCs) and EpiSC from mouse embryonic fibroblasts (MEFs). (**C**) Brightfield images of F1 hybrid-derived colonies of iPSCs (top) and EpiSCs (bottom). Scale bar = 100μm. (**D**) Principal component analysis (PCA) on bulk RNA-seq datasets from F1-hybrid EpiSCs. (**E**) Immunostaining in iPSCs and EpiSCs, from the SpretB6 background, against the general pluripotency markers NANOG, SOX2, and OCT4, and the naïve pluripotency marker KLF4. Scale bar = 30μm. (**F**) Heatmap representation of gene expression as detected by bulk RNA-seq on F1-hybrid EpiSCs. The name of the cell line indicates the background and the clone number. (**G**) Differentiation protocol and immunostaining of EpiSC-derived mesoderm cells against the mesoderm marker T and the paraxial mesoderm marker TBX6. Scale bar = 100μm. (**H**) Differentiation protocol and immunostaining of EpiSC-derived definitive endoderm cells against the endoderm markers SOX17 and FOXA2. Scale bar = 100μm.

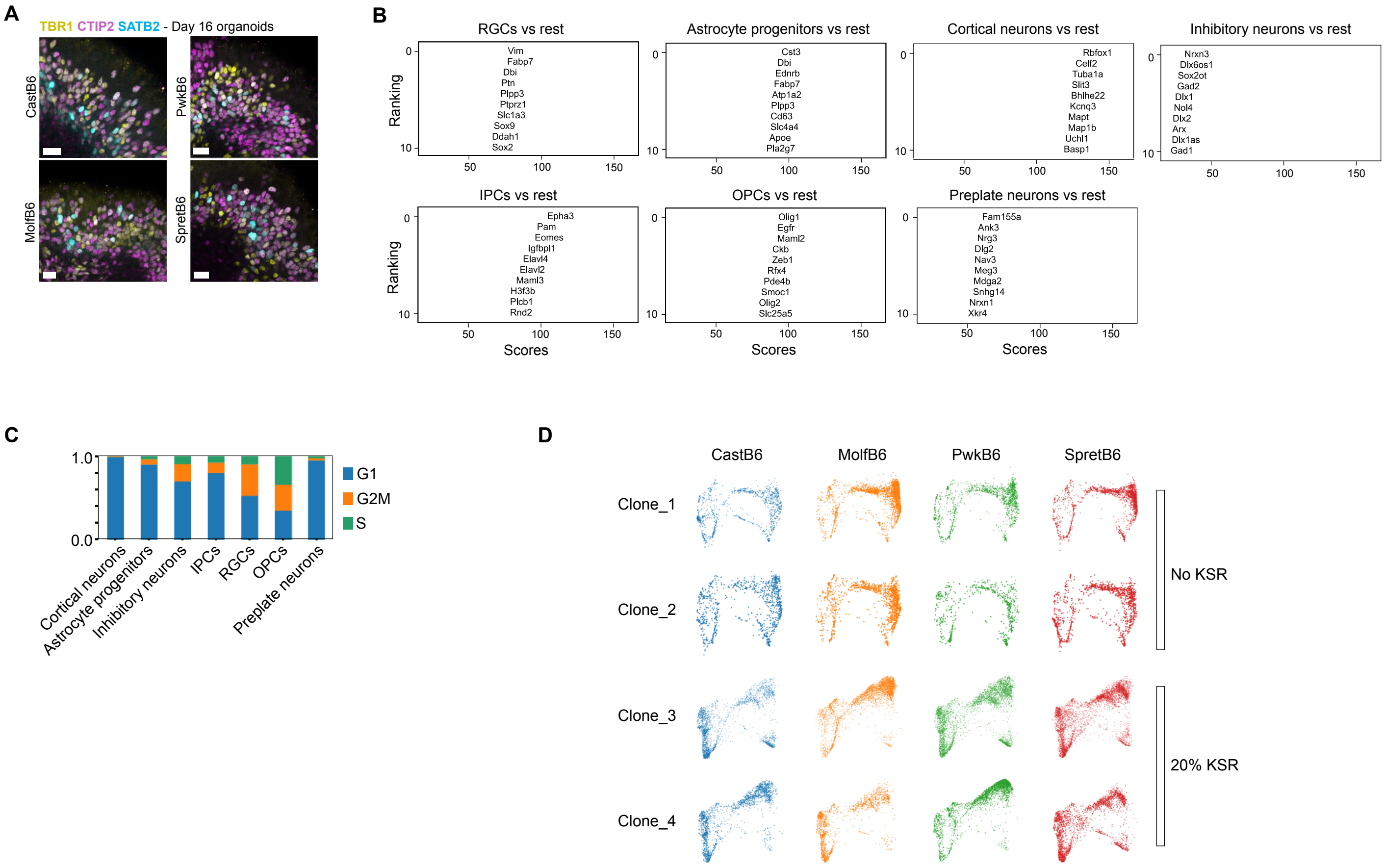


**Supplementary figure 3.** Related to main figure 5. (**A**) Immunostaining of d16 neocortical organoids from F1-hybrid EpiSCs for markers of deep-layer cortical neurons (TBR1 and CTIP2) and upper-layer cortical neurons (SATB2). Scale bar = 20µm. (**B**) Wilcoxon rank-sum test across all cell types detected by scRNA-seq on day 12 organoids, the top 10 ranked genes are displayed. (**C**) Cell cycle analyses on the cell types detected by scRNA-seq on d12 organoids. (**D**) Force-directed layout representation of each clone used for scRNA-seq (d12), separated by whether they were treated with 20% KSR from d4 or not.

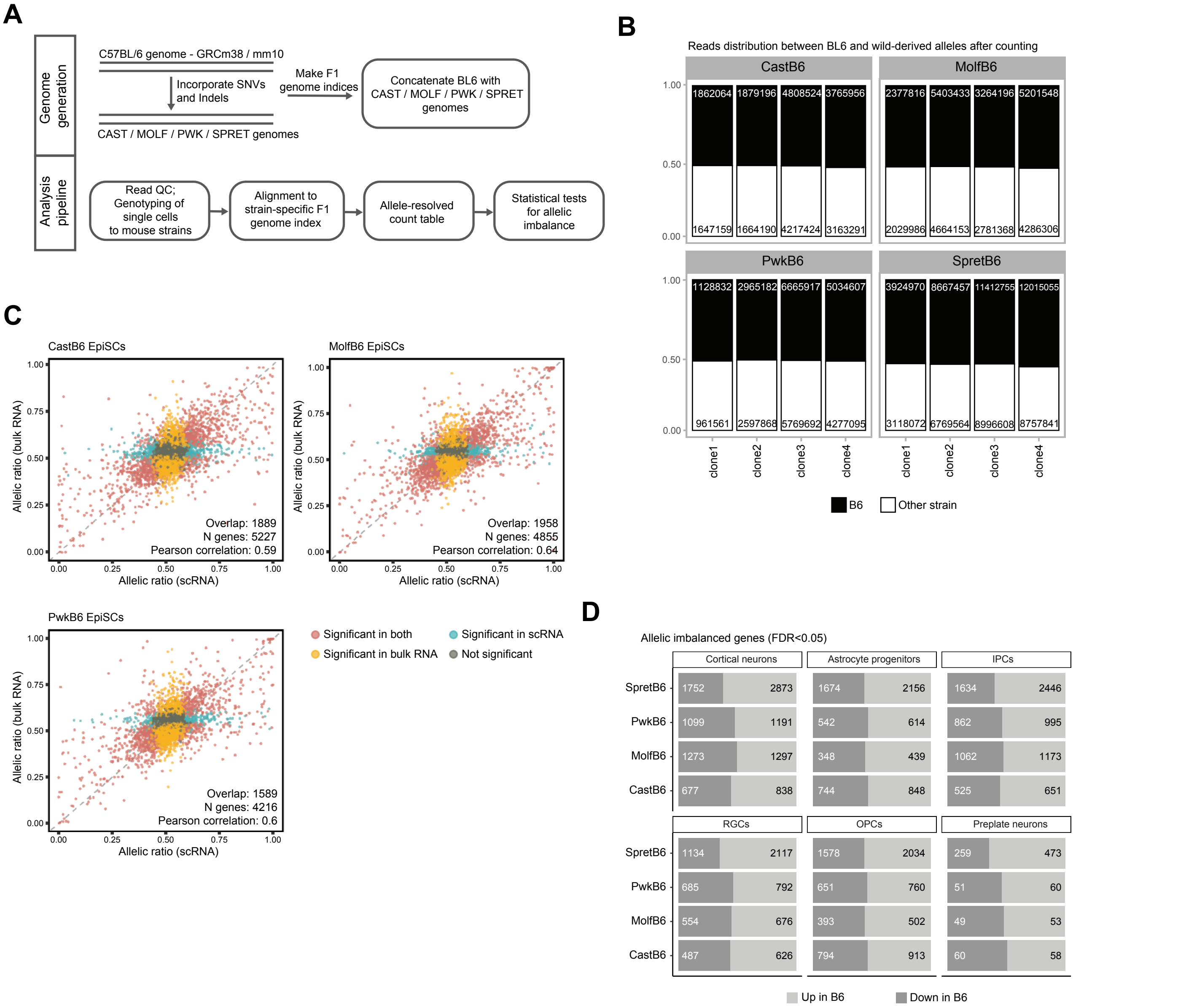


**Supplementary figure 4.** Related to main figure 6. (**A**) General pipeline for ASPEN, our new method for allelic mapping in single cells. (**B**) Bar plots of the number of reads, as detected by scRNA-seq on d12 organoids, mapped to the B6 or the wild-derived genome across clones and backgrounds. (**C**) Correlation of allelic ratios between bulk RNA-seq and scRNA-seq on CastB6, MolfB6, and PwkB6 EpiSCs, collected from two independent batches. The Pearson coefficient and the number of genes showing overlapping behavior in both datasets are displayed. (**D**) Allelic imbalances per cell type and background subdivided by whether they are skewed towards the B6 or the wild-derived allele.

**A**

| | CastB6 | MolfB6 | PwkB6 | SpretB6 |
|---|---|---|---|---|
| Preplate neurons | 0.27 | 0.30 | 0.22 | 0.20 |
| OPCs | 0.23 | 0.22 | 0.24 | 0.39 |
| RGCs | 0.17 | 0.21 | 0.24 | 0.38 |
| IPCs | 0.19 | 0.29 | 0.31 | 0.40 |
| Astrocyte progenitors | 0.18 | 0.17 | 0.17 | 0.34 |
| Cortical neurons | 0.24 | 0.38 | 0.35 | 0.45 |

**B**

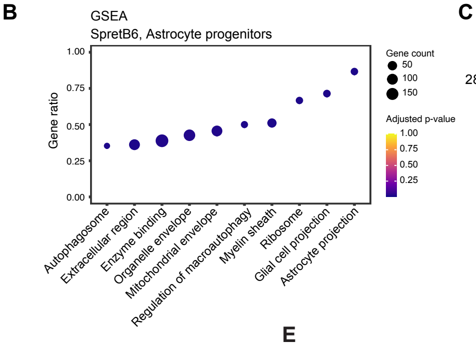


**C**

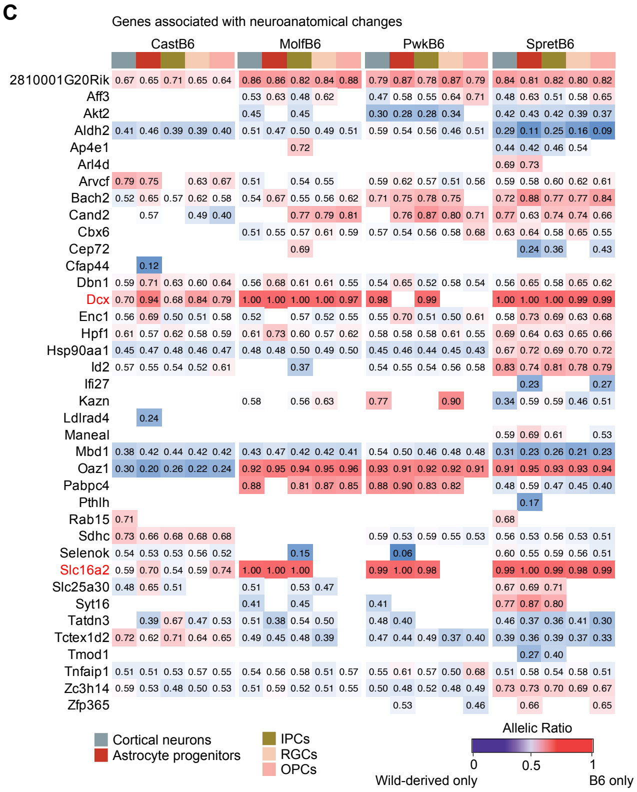


**D**

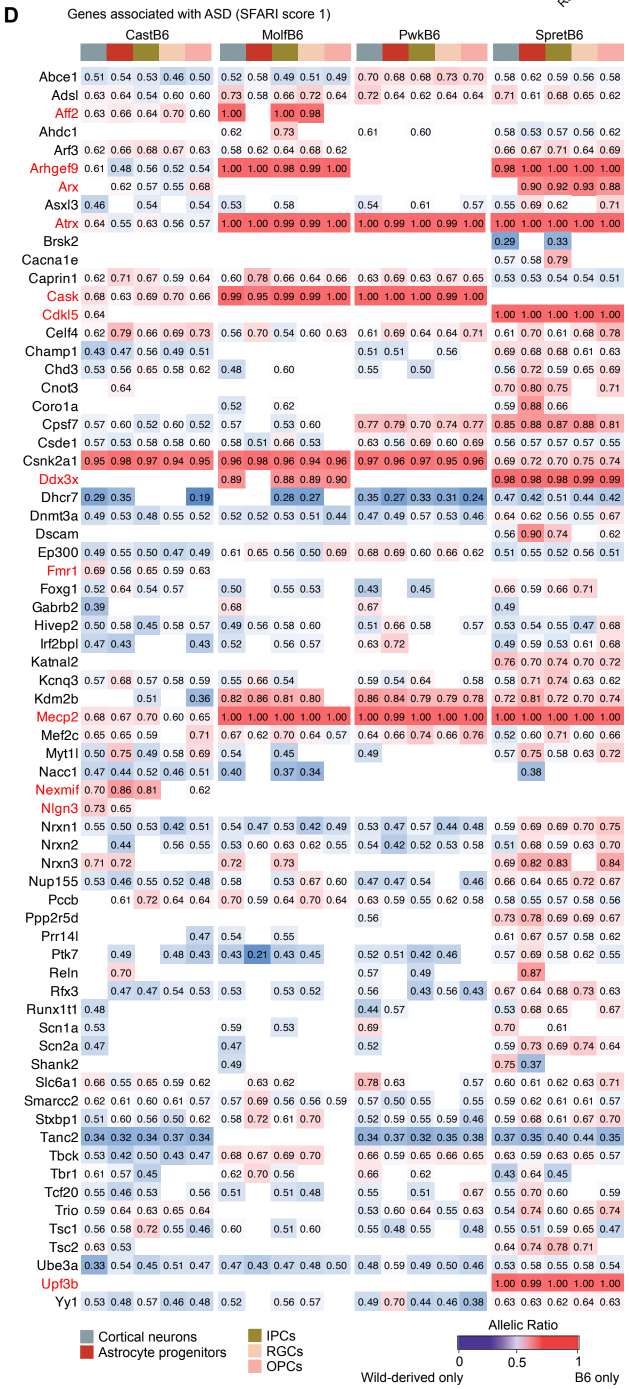


**E**

Total genes:1,688
Genes with BIC diff >= 10: 718

| Model | n | freq |
|---|---|---|
| Model1 | 109 | 0.15 |
| Model2 | 340 | 0.47 |
| Model3 | 219 | 0.31 |
| Model4 | 50 | 0.07 |
| Total | 718 | 1 |

**F**

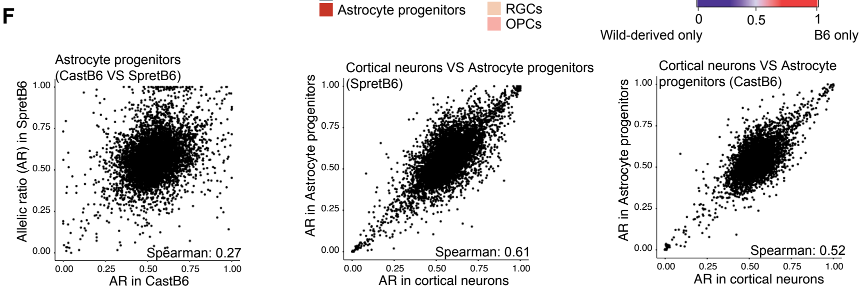


**G**

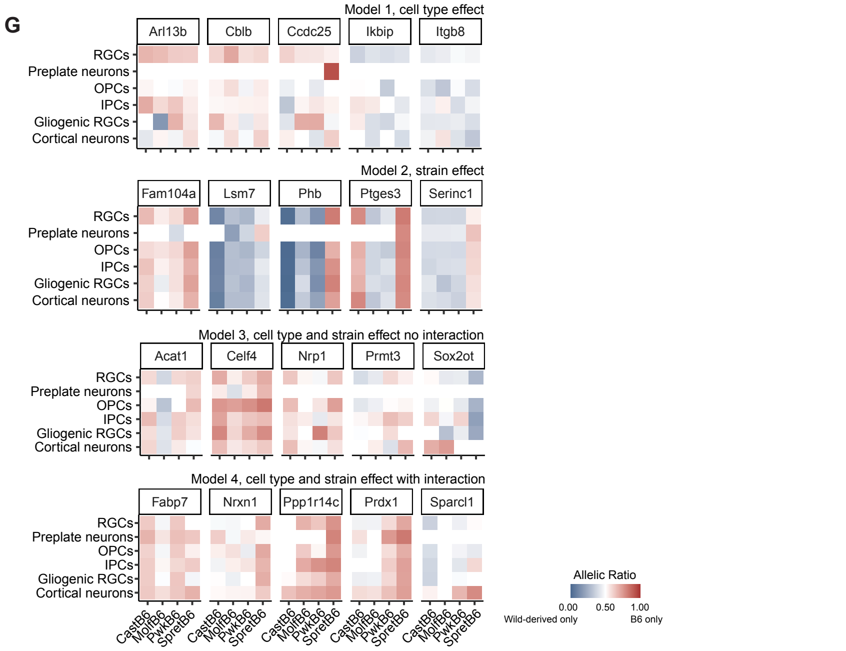

**Supplementary figure 5.** Related to main figure 6. (**A**) Proportion of genes detected as AI across genetic backgrounds and cell types. (**B**) GSEA analyses on the AI genes in SpretB6 astrocyte progenitors. (**C**) Allelic ratio per cell type and background for the AI genes linked to neuroanatomical changes by the International Mouse Phenotyping Consortium 31. Genes in red are located in the X-chromosome. (**D**) Allelic ratio per cell type and background for the AI genes associated with ASD (SFARI database, score 1 only). Genes in red are located in the X-chromosome. (**E**) The number of genes found to have a better fit in of the four generalized linear mixed models, based on BIC > 10 between the two competing models. Allelic ratio variance was better explained by cell type (Model 1), by genetic background (Model 2), by genetic background and cell type jointly (Model 3), by an interaction between genetic background and cell type (Model 4). (**F**) Correlation of allelic ratios across strains and cell types. The Spearman correlation value is displayed. (**G**) Allelic expression across four generalized linear mixed models, highlighting the differences due to either cell type or genetic background effects or a combination of both.

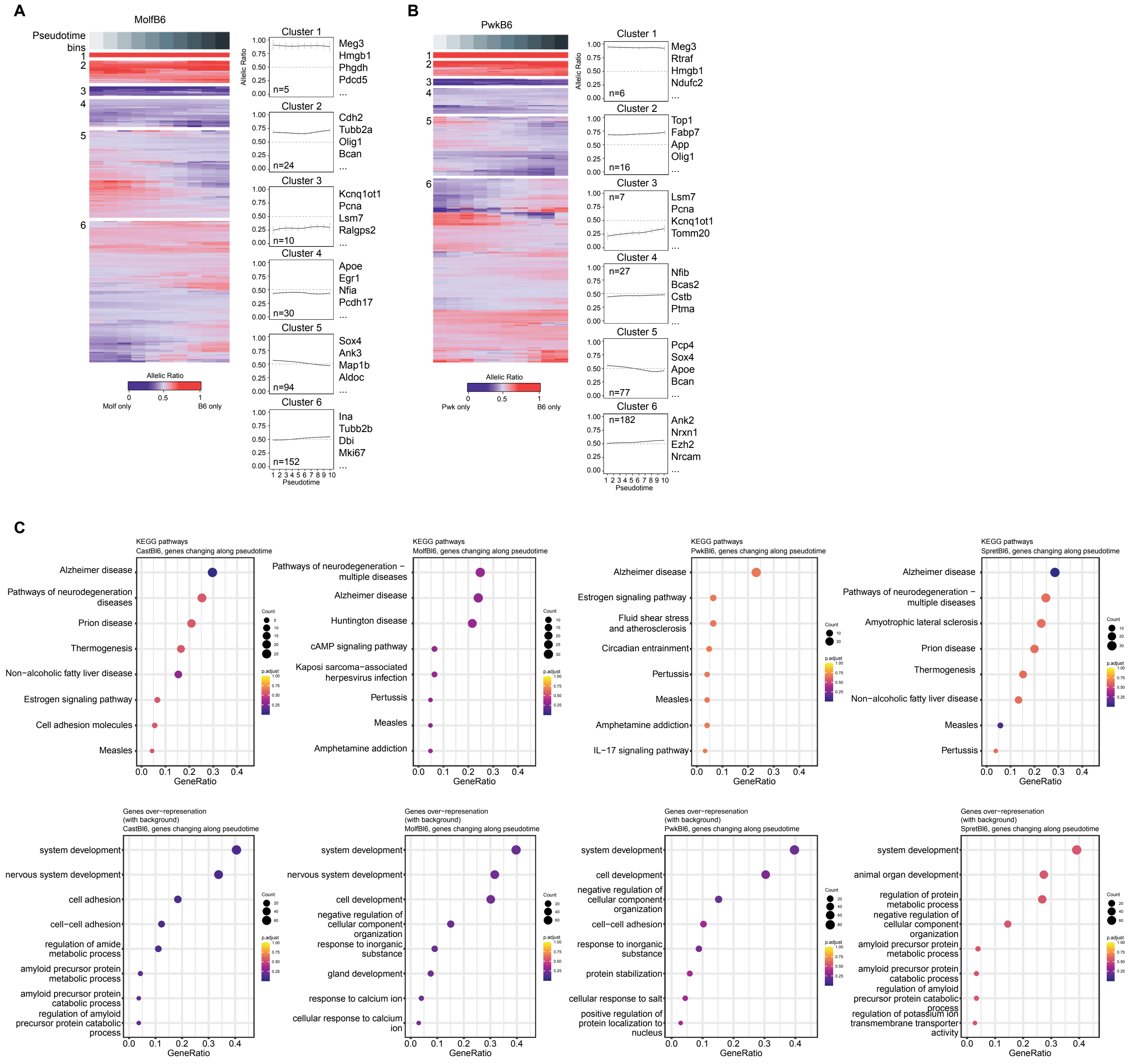


**Supplementary figure 6.** Related to main figure 7. (**A**) Heatmap of allelic ratio trends along the neurogenic pseudotime, for MolfB6 and (**B**) PwkB6 organoids, for 315 genes with significant association (GAM model fit) in at least one background, grouped using hierarchical clustering. Values are averaged within 10 pseudotime bins. Line plots (right) show individual gene trajectories in grey and the average trend in black for each cluster. (**C**) KEGG pathway enrichment analyses (top) and over-representation analyses (bottom) on genes within clusters 5 and 6 (e.g., genes that change behavior along pseudotime) across backgrounds.
