## Supplementary material for "A MOUSE ORGANOID PLATFORM FOR MODELING CEREBRAL CORTEX DEVELOPMENT AND CIS-REGULATORY EVOLUTION IN VITRO": Key resources table

| REAGENT or RESOURCE | SOURCE | IDENTIFIER |
| --- | --- | --- |
| Antibodies |  |  |
| Rabbit pAb anti-GFP | Abcam | Cat#AB290; RRID:AB_303395 |
| Goat pAb anti-KLF4 | R&D Systems | Cat#AF3158; RRID:AB_2130245 |
| Rabbit mAb anti-NANOG (D2A3) | Cell Signaling Technology | Cat#8822S; RRID:AB_11217637 |
| Mouse mAb anti-OCT3/4 (C10) | Santa Cruz Biotechnology | Cat#SC-5279; RRID:AB_628051 |
| Rabbit pAb anti-SOX2 | Millipore | Cat#AB5603; RRID:AB_2286686 |
| Goat pAb anti-TBX6 | R&D Systems | Cat#AF4744; RRID:AB_2200834 |
| Rabbit mAb anti-T/Brachyury | Abcam | Cat#AB209665;RRID:AB_2750925 |
| Goat pAb anti-SOX17 | R&D Systems | Cat#AF1924; RRID:AB_355060 |
| Rabbit mAb anti-FOXA2 | Abcam | Cat#AB108422; RRID:AB_11157157 |
| Rabbit pAb anti-BF1/FOXG1 | Takara | Cat#M227; RRID:AB_2827749 |
| Rabbit pAb anti-TBR1 | Abcam | Cat#ab31940; RRID:AB_2200219 |
| Chicken pAb anti-TBR2 | Millipore | Cat#AB15894; RRID:AB_10615604 |
| Rat mAb anti-GFAP (2.2B10) | Thermo Fisher Scientific | Cat#13-0300; RRID:AB_2532994 |
| sdAb anti-GFAP - conjugated with FluoTag-X2 | NanoTag Biotechnologies | Cat#N3802-AF568-L; RRID: AB_3076115 |
| Rat mAb anti-CTIP2 (25B6) | Abcam | Cat#AB18465; RRID:AB_2064130 |
| Mouse mAb anti-CUX1/CUTL1/CASP (2A10) | Abcam | Cat#ab54583; RRID:AB_941209 |
| Mouse mAb anti-PAX6 | BD Biosciences | Cat#561462; RRID:AB_10715442 |
| Mouse mAb anti-SATB2 (SATBA4B10) | Abcam | Cat#AB51502; RRID:AB_882455 |
| Chicken pAb anti-NESTIN | Aves Labs | Cat#NES; RRID:AB_2314882 |
| Rabbit mAb anti-OLIG2 (EPR2673) | Abcam | Cat#AB109186; RRID:AB_10861310 |
| Goat pAb anti-POU3F3/BRN1 | Novus Biologicals | Cat#NBP1-49872; RRID:AB_10012062 |
| Rat mAb anti-MBP (Clone 12) | Novus Biologicals | Cat#NB600-717; RRID:AB_2139899 |
| Rabbit mAb anti-S100β (EP1576Y) | Abcam | Cat#AB52642; RRID:AB_882426 |
| Goat pAb anti-PDGFRα | R&D Systems | Cat#AF1062; RRID:AB_2236897 |
| Rabbit mAb anti-NEUN (EPR12763) | Abcam | Cat#AB209898; RRID:AB_2532109 |
| Mouse mAb anti-CNP (CL2887) | Atlas Antibodies | Cat#AMAB91072; RRID:AB_2665789 |
| Chicken pAb anti-NF-H (Poly28226) | BioLegend | Cat#822601; RRID:AB_2564859 |
| Alexa Flour 488 Donkey anti-Rat | Thermo Fisher Scientific | Cat#A21208; RRID:AB_2535794 |
| Alexa Flour 647 Donkey anti-Rat | Thermo Fisher Scientific | Cat#A48272TR; RRID:AB_2896338 |
| Alexa Flour 488 Donkey anti-Rabbit | Thermo Fisher Scientific | Cat#A21206; RRID:AB_2535792 |
| Alexa Flour 555 Donkey anti-Rabbit | Thermo Fisher Scientific | Cat#A31572; RRID:AB_162543 |
| Alexa Flour 647 Donkey anti-Rabbit | Thermo Fisher Scientific | Cat#A31573; RRID:AB_2536183 |
| Alexa Flour 555 Donkey anti-Mouse | Thermo Fisher Scientific | Cat#A31570; RRID:AB_2536180 |
| Alexa Flour 647 Donkey anti-Mouse | Thermo Fisher Scientific | Cat#A31571; RRID:AB_162542 |
| Alexa Flour 488 Donkey anti-Goat | Thermo Fisher Scientific | Cat#A32814; RRID:AB_2762838 |
| Alexa Flour 555 Donkey anti-Goat | Thermo Fisher Scientific | Cat#A21432; RRID:AB_2535853 |
| Alexa Flour 647 Donkey anti-Goat | Thermo Fisher Scientific | Cat#A32849; RRID:AB_2762840 |
| Alexa Flour 555 Goat anti-Chicken | Thermo Fisher Scientific | Cat#A21437; RRID:AB_2535858 |
| Alexa Flour 647 Donkey anti-Chicken | Jackson ImmunoResearch | Cat#703605155; RRID:AB_2340379 |
| Plasmid and virus strains |  |  |
| CytoTune emGFP Sendai fluorescence reporter | ThermoScientific | Cat#A16519 |
| TetO-FUW-OSKM | Addgene | RRID:Addgene_20321 |
| FUW-M2rtTA | Addgene | RRID:Addgene_20342 |
| Chemicals, peptides, and recombinant proteins |  |  |
| 2-Mercaptoethanol | Gibco | Cat#21985023 |
| 16% Formaldehyde (w/v), Methanol free | Thermo Scientific | Cat#28908 |
| Accutase | Gibco | Cat#A1110501 |
| Activin A | Peprotech | Cat#120-14P |
| Anti-adherence rinsing solution | STEMCELL Technologies | Cat#7010 |
| Apo-transferrin | InVitra | Cat#777TRF029 |
| B27 supplement | ThermoScientific | Cat#17504044 |
| *Continued* |  |  |
| REAGENT or RESOURCE | SOURCE | IDENTIFIER |
| B27 supplement (without vitamin A) | Gibco | Cat#12587010 |
| BenchMark Fetal bovine serum | Gemini | Cat#100-106 |
| Bovine Serum Albumin | Sigma | Cat#A2153 |
| Cell recovery solution | Corning | Cat#354253 |
| Chemically Defined Lipid Concentrate | Gibco | Cat#11905031 |
| CHIR99201 | Tocris Bioscience | Cat#252917-06-9 |
| Chroman 1 | MedChem Express | Cat#HY-15392 |
| Collagenase Type IV | Gibco | Cat#17104019 |
| Cyclopamine | Cayman Chemical | Cat#11321 |
| DeepClear | Celexplorer | Cat#DC-201 |
| Dimethyl sulfoxide (DMSO) | Sigma-Aldrich | Cat#D2650 |
| DMEM with glutamax | Gibco | Cat#10564029 |
| DMEM/F12 medium | Gibco | Cat#11320082 |
| DNase I | Zymo Research | Cat#E1010 |
| Doxycycline hyclate | Sigma-Aldrich | Cat#D9891-1G |
| EDTA (0.5 M), pH 8.0, RNase-free | Invitrogen | Cat#AM9261 |
| Emricasan | Selleck Chemicals | Cat#S7775 |
| ESGRO LIF | Sigma | Cat#ESG1107 |
| F12 with GlutaMAX | Gibco | Cat#31765035 |
| Fibronectin | Sigma | Cat#FC010 |
| Glutamax | Gibco | Cat#35050061 |
| Heat stable recombinant human bFGF | Gibco | Cat#PHG0360 |
| Hyclone fetal bovince serum | Cytiva | Cat#SH3007003 |
| IMDM | Gibco | Cat#12440053 |
| Insulin | Sigma | Cat#91077C |
| KaryoMAX Colcemid | Thermo Fisher Scientific | Cat#151212012 |
| Knockout serum replacement | Gibco | Cat#10828028 |
| Laminin from EHS murine sarcoma basement membrane | Sigma-Aldrich | Cat#L2020 |
| L-Ascorbic Acid | FUJIFILM Wako Chemicals | Cat#32344822 |
| LDN-193189 | Stemgent | Cat#04-0074 |
| LGK-974 | Selleck Chemicals | Cat#S7143 |
| LY-294002 | Selleck Chemicals | Cat#S1105 |
| Matrigel | Corning | Cat#354230 |
| Monothioglycerol | Sigma | Cat#M6145 |
| N2 supplement | Gibco | Cat#17502048 |
| Neurobasal Media | Gibco | Cat#21103049 |
| Non-essential amino acids | Gibco | Cat#11140050 |
| NVP-TNKS656 | Selleck Chemicals | Cat#S7238 |
| PD0325901 | Tocris | Cat#4192 |
| Penicillin/streptomycin (100X) | Gibco | Cat#15140122 |
| Polybrene (Hexadimethrine bromide) | Sigma-Aldrich | Cat#H9268 |
| Polyvinyl alcohol | Sigma-Aldrich | Cat#341584 |
| Potassium chloride solution | Sigma-Aldrich | Cat#P9327 |
| rhLaminin-521 | Gibco | Cat#A29249 |
| SB-431542 | Tocris | Cat#1614 |
| Sodium Dodecyl Sulfate (SDS) | Fisher Scientific | Cat#BP24361 |
| Sodium pyruvate | Gibco | Cat#11360070 |
| TRI Reagent | Sigma-Aldrich | Cat#T9424 |
| Triton X-100 | Sigma-Aldrich | Cat#9002-93-1 |
| TrypLE | Thermo Scientific | Cat#12605028 |
| VECTASHIELD Antifade mounting medium with DAPI | Vector Laboratories | Cat#H1200 |
| *Continued* |  |  |
| REAGENT or RESOURCE | SOURCE | IDENTIFIER |
| Critical commercial assays |  |  |
| PCR Mycoplasma Detection Kit | ABM | Cat#G238 |
| DNA Clean and Concentrator Kit | Zymo Research | Cat#D4014 |
| Chromium Next GEM Single Cell 3' Kit v3.1, 16 rxns | 10x Genomics | Cat#1000406 |
| Chromium Next GEM Single Cell ATAC Kit v2, 4 rxn | 10x Genomics | Cat#1000406 |
| Experimental models: cell lines |  |  |
| Listed in Supplemental Table 1 |  |  |
| Software and algorithms |  |  |
| Adobe Illustrator | Adobe | https://www.adobe.com/ |
| R studio | R studio development team | https://www.rstudio.com/ |
| Imaris Viewer 10.1.1 | Oxford Instruments | https://imaris.oxinst.com/imaris-viewer |
| Scanpy (Python package) | scverse project | https://scanpy.readthedocs.io |
| Deposited data |  |  |
| scRNA-seq data from Organoids Day12 | This paper | GSE268332 |
| scRNA-seq data from EpiSCs | This paper | GSE268332 |
| Bulk RNA-seq data from EpiSCs | This paper | GSE268329 |
| snRNA-seq from Organoids Day18 | This paper | GSE268332 |
| scATAC-seq data from Organoids Day18 | This paper | GSE268331 |
| Other |  |  |
| AggreWell 400, 24-well plate | STEMCELL Technologies | Cat#34411 |
| 24-well suspension culture plate | CELLSTAR | Cat#662102 |
| Superfrost plus microscope slides, white tab | Fisher Scientific | Cat#1255015 |
| Microscope cover glass (24x50mm) | Globe Scientific Inc. | Cat#1415-10 |
| Kord Agri-plate petri dish (100mmx25mm) | Fisher Scientific | Cat#2906 |
| 6-well ultra-low adherent plate | Corning | Cat#3471 |
| Leica SP8 confocal microscope | Leica Microsystems | N/A |
