## Supplementary experimental methods for "A MOUSE ORGANOID PLATFORM FOR MODELING CEREBRAL CORTEX DEVELOPMENT AND CIS-REGULATORY EVOLUTION IN VITRO"

**Protocol: Generation of Cortical Organoids from Mouse EpiSCs**

Important notes before starting:

- Before use, all media should be warmed to 37°C using a water or bead bath.
- All steps should be performed in a sterile BSC using rigorous aseptic technique.

Formation of EBs

1. Confirm EpiSC quality (approximately 80% confluency) using a phase contrast microscope.
2. Aspirate EpiSC media from the plate.
3. Using a serological pipette, wash each well/plate twice with PBS ^-/-^.
4. Aspirate PBS ^-/-^ from plate and apply prewarmed 0.5 µg/µL Collagenase IV to each well/plate to preferentially dissociate EpiSC colonies without feeder contamination.
   a. *Optional:*
      i. *After applying Collagenase IV, place the plate back into the incubator for no more than 5 minutes to increase the speed of dissociation.*
      ii. *Gently tap the sides of the plate to encourage colony lifting.*
      iii. *Do not leave the Collagenase IV longer than 25-30 minutes at room temperature as this will cause feeder cells to dissociate as well.*
5. Once the EpiSC colonies are lifted, gently collect colonies using a serological pipette and transfer into a 15 mL centrifuge tube.
6. Add an equal volume of HBS++ to the same 15 mL centrifuge tube to dilute the Collagenase IV.
   a. *Optional:*
      i. *Wash each plate gently with HBS++ to collect any remaining colonies and combine with the Collagenase IV and PBS ^-/-^ in the 15 mL tube.*
7. Centrifuge the cells at 200 g for 3 minutes to pellet EpiSC colonies and aspirate the supernatant.
8. Using a serological pipette, resuspend the pellet with 1 mL of Accutase to break colonies into a single-cell suspension. Leave the cell suspension in the hood for 2 minutes to ensure single-cell suspension.
   a. *Optional:*
      i. *Remove one drop of the suspension (~10 µL) and place it on a microscope slide. Observe under the microscope for residual colonies. If there are residual colonies present, the cell suspension may require no more than 2 more minutes under the hood to acquire single-cell suspension. Continue to observe the cell suspension under the microscope during this time to ensure single-cell suspension.*
      ii. *Gently pipette the suspension using a serological pipette once the cell suspension has been in the hood for at least 1 minute to ensure single-cell suspension.*
9. Once the cells are completely dissociated, add 5-10 mL of PBS ^-/-^ to dilute the Accutase.
10. Centrifuge the solution at 200 g for 3 minutes to pellet the single cells.
11. To prepare Aggrewell plate, open it in the hood and add 500 µL of anti-adherence rinsing solution to each well, pipette up and down a few times gently.
12. Centrifuge the plate at 1300 g for 5 minutes along with an identical balance Aggrewell plate with 500 µL of PBS ^-/-^ in each well. Aspirate anti-adherence solution.
13. Rinse each Aggrewell well with 2 ml of EB formation media (Table 1) and aspirate. Load 1mL of EB formation media in each well and place Aggrewell plate in incubator.
14. Aspirate the supernatant, resuspend in the EB formation media and count the cells.
15. For EBs of 1000 cells each, calculate the appropriate volume of media and the number of cells to seed the desired number of wells, then prepare the master solution with 1,200,000 cells/well of an Aggrewell plate, with each well containing a total of 2 mL of media.
16. Using a serological pipette, disperse the master solution into well(s) of the Aggrewell plate.
17. Using a P1000 pipette, gently pipet up and down to equally distribute the cells within the well(s).
18. Centrifuge the Aggrewell plate with a symmetrical balance at 100 g for 3 minutes. Confirm cells are evenly distributed using a microscope.
19. Place Aggrewell in an incubator for 24 hours to form EBs. Avoid moving the plate during the EB formation phase.

Collection and Embedding of Organoids

Important note before starting:

- Matrigel should be thawed overnight on ice at 4°C or for at least 3-4 hours before collecting and embedding organoids

1. Following 24 hours in EB formation media, the Aggrewell should contain uniform, well-formed developing organoids. Appearance of organoids can be confirmed using a phase contrast microscope. Using a P1000 pipette, gently remove as much media as possible from the Aggrewell without disturbing the organoids.
2. Using a P1000 with a wide orifice tip, add 1 mL of Neural Induction media (Table 1) to the Aggrewell and gently resuspend the developing organoids.
3. Prepare 1 6 cm Petri dish with 5 ml of Neural Induction media and 1 10 cm Petri dish with 12.5 ml of Neural Induction media for each well of the Aggrewell plate.
4. Using a wide-orifice tip, transfer organoids to the 6 cm Petri dish containing 5 mL of Neural Induction media and swirl briefly to dilute the remaining EB formation media. Transfer organoids to the 10 cm Petri dish containing 10 mL of Neural Induction media.
   a. *Note: Washing with PBS ^-/-^ should be avoided as this may cause organoids to adhere to the plate.*
5. Determine the total number of organoids desired and how many wells of a 6 well plate are needed.
   a. *Note: Each well of the plate should contain 20-100 organoids.*
6. Carefully gather 20-100 organoids in 67 µL of Neural Induction media and pipette up and down to mix with 100 µL of Matrigel in an Eppendorf tube. Carefully dispense the mix of Matrigel, media, and organoids (167 µL) in the center of a well of the 6 well plate, being sure to avoid the walls of the well (Derived from Qian et al., 2018).
7. Repeat for desired number of wells and then place the plate in the incubator for 30 minutes to solidify the Matrigel dome.
8. Following the 30-minute incubation period, carefully remove the plate from the incubator and gently add 3 mL of Neural Induction media to each well, pipetting onto the sides of the well to avoid disturbing the Matrigel dome. Return to the incubator for 24 hours.
   a. *Note: When adding the Neural Induction media, the Matrigel dome might float off the plate. This does not affect the quality of the organoids.*

Maintenance and Patterning of Organoids

1. Following 24 hours in Neural Induction media, gently remove as much media as possible without disturbing the Matrigel domes.
   a. *Note: Wash and media change should both be performed under a dissection microscope to avoid organoid loss.*
2. Add 3 mL of Neuroepithelial Expansion media (Table 1) to each well, pipetting onto the sides of the well. Place plate in an incubator for 24 hours.
3. Following 24 hours in Neuroepithelial Expansion media, gently remove as much media as possible without disturbing the Matrigel domes.
4. Add 3 mL of Wnt Activated Neuroepithelial Expansion media (Table 1) to each well, pipetting onto the sides of the well. Place plate in an incubator for 24 hours.
5. After 24 hours in the Wnt Activated Neuroepithelial Expansion media, gently remove the media and wash twice with PBS ^-/-^.
6. After removing the second PBS ^-/-^ wash, add 2 mL of Corning Cell Recovery Solution to each well of the 6-well plate, ensuring the domes are lifted from the plastic surface. Gently swirl the plate and place at 4°C for 30 minutes to remove Matrigel from organoids.
   a. *Optional:*
      i. *To improve recovery, gently swirl plate every 10 minutes while at 4°C.*
7. While organoids are at 4°C for Matrigel removal, prepare 1 6 cm Petri dish with 5 ml of Neuronal media and 1 10 cm Petri dish with 12.5 ml of Neuronal media for each well of the 6 well plate containing organoids.
8. Following Matrigel removal, gently add 2 mL of Neuronal media (Table 1) to each well of the 6-well plate to dilute the solution.
9. Using a wide orifice tip, transfer the organoids from a well of a 6-well plate to the 6 cm Petri dish containing 5 mL of Neuronal media to ensure maximum dilution of the remaining Corning Cell Recovery solution.
10. Using a wide orifice tip, transfer organoids from the 6 cm Petri dish to the 10 cm Petri dish containing 12.5 mL of Neuronal media. Place an empty shallow 10 cm Petri dish on the shaker and place the 10 cm Petri dish containing the organoids on top to avoid direct contact with the metal.
    a. *Note: The speed of the shaker will depend on the specific shaker being used. For Celltron benchtop shaker, 65 rpm is sufficient to prevent organoid merging.*
11. Change media using fresh Neuronal media every other day until collection.

**Protocol: Immunostaining of Mouse EpiSC-derived Cortical Organoids (method derived from Dekkers et al., 2019)**

Fixation and Immunostaining of Organoids

1. Coat a 15 mL centrifuge tube with 1% BSA in PBS ^-/-^ to avoid organoid attachment to the tube and place on ice.
2. Collect organoids using a wide orifice tip and, while on ice, gently add organoids to the pre-coated centrifuge tube and wash three times with 4°C PBS ^-/-^.
3. To fix organoids, add 4% PFA in PBS ^-/-^ to 15 mL centrifuge tube until organoids are completely submerged. Place on ice for 45 minutes and gently resuspend halfway through incubation to ensure proper fixation.
   a. *Note: When fixing organoids aged 2 months or older, proper fixation and immunostaining can be achieved by cutting organoids in half with a sterile razor blade prior to fixation.*
4. Following fixation, wash three times for 5 minutes each with PBS ^-/-^ containing 0.1% Triton-X (PBST) to remove any remaining PFA.
5. To permeabilize organoids, add PBS ^-/-^ containing 0.5% Triton-X until organoids are completely submerged and incubate at 4°C.
   a. *Note: Incubation time varies based on age and size of organoids. Typically, Day 4 or younger organoids are incubated for ~15 minutes, Day 5-16 organoids are incubated for ~30 minutes, Day 16 and older organoids are incubated for ~45 minutes.*
6. Following incubation period, remove permeabilization buffer and block organoids for 15 minutes in organoid washing buffer (OWB) containing 0.2% Triton-X, 0.02% SDS, and 0.2% BSA in PBS ^-/-^. Calculate appropriate volume of OWB by determining number of staining conditions (200 µL of OWB per condition).
7. Equally distribute the organoids in a low-adhesion 24-well plate with a total of 200 µL OWB and desired number of organoids in each well. Each well will represent a single staining condition.
8. Following 15-minute blocking period, directly add primary antibodies of interest prediluted in 200 µL OWB to each well for a total volume of 400 µL per well.
9. Place on shaker and incubate primary antibodies overnight at 4°C.
   a. *Note: Primary antibody incubation can last between an overnight period and up to 2 weeks long.*
10. Following overnight incubation, add 800 µL of OWB directly to each well and place on shaker at room temperature for 5 minutes.
11. Next, remove 1 mL of solution, replace with fresh 1 mL of OWB, and place on shaker for 30 minutes at room temperature to wash organoids.
12. Repeat this wash step at least two additional times, for a minimum total of 1.5 hours of washing.
13. After at least 1.5 hours of washing, remove 1 mL of OWB from each well and directly add 200 µL of prediluted secondary antibodies and DAPI (final concentration of 1 µM) in OWB, for a total volume of 400 µL per well.
14. Place on shaker and incubate for at least 24 hours at 4°C.
15. Following secondary antibody incubation, remove from 4°C, and add 800 µL of OWB per well. Place on shaker at room temperature for 5 minutes.
16. Next, remove 1 mL of OWB from each well and replace with a fresh 1 mL of OWB. Place on shaker at room temperature for 30 minutes to wash organoids.
17. Repeat this step at least two additional times, for a minimum total of 1.5 hours of washing.

Preparation of Slide for Imaging

1. Using a standard hole puncher, punch a hole (0.5 cm Ø) into a sticky silicone pad (20x20x0.5 mm).
2. Place sticky pad on microscope slide (Figure 1)

Mounting of Organoids for Imaging

1. Using a wide orifice tip, transfer organoids from one well of Aggrewell plate to a 1.5 mL Eppendorf tube.
2. Remove as much supernatant as possible without disturbing the organoids.
   a. *Note: If the organoids are at Day 4 or less, centrifuge at 70 g for 3 minutes to pellet.*
3. Using a wide orifice pipette tip, add ~ 40 µL of DeepClear solution and gently resuspend the organoids. Transfer the organoid resuspension to a prepared slide. Add coverslip and image.

**Table 1.** Media Composition for Neural Organoids

| EB Formation media | Neural Induction media | Neuroepithelial Expansion media | Wnt Activated Neuroepithelial Expansion media | Neuronal media |
| --- | --- | --- | --- | --- |
| N2B27 (B27 without vitamin A) | N2B27 (B27 without vitamin A) | N2B27 (B27 without vitamin A) | N2B27 (B27 without vitamin A) | N2B27 (B27 without vitamin A) |
| 50 nM chroman-1 | 100 nM LDN | 1% KSR | 1% KSR | 20% KSR |
| 5 µM emricasan | 10 µM SB431542 | 1 µM cyclopamine | 1 µM cyclopamine |  |
| 100 nM LDN | 100 nM LGK974 |  | 3 µM CHIR |  |
| 10 µM SB431542 | 1 µM cyclopamine |  |  |  |
| 100 nM LGK974 |  |  |  |  |
| 1 µM cyclopamine |  |  |  |  |

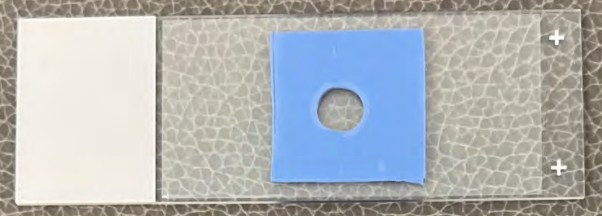

**Figure 1.** Slide for Organoid Imaging
